# A post-injury immune challenge with lipopolysaccharide following adult traumatic brain injury alters neuroinflammation and the gut microbiome acutely, but has little effect on chronic outcomes

**DOI:** 10.1101/2024.09.28.615631

**Authors:** Sarah S.J. Rewell, Ali Shad, Lingjun Chen, Matthew Macowan, Erskine Chu, Natasha Gandasasmita, Pablo M. Casillas-Espinosa, Jian Li, Terence J. O’Brien, Bridgette D. Semple

## Abstract

Patients with a traumatic brain injury (TBI) are susceptible to hospital-acquired infections, presenting a significant challenge to an already-compromised immune system. The consequences and mechanisms by which this dual insult worsens outcomes are poorly understood. This study aimed to explore how a systemic immune stimulus (lipopolysaccharide, LPS) influences outcomes following experimental TBI in young adult mice. Male and female C57Bl/6J mice underwent controlled cortical impact or sham surgery, followed by 1 mg/kg i.p. LPS or saline-vehicle at 4 days post-TBI, before behavioral assessment and tissue collection at 6 h, 24 h, 7 days or 6 months. LPS induced acute sickness behaviors including weight loss, transient hypoactivity, and increased anxiety-like behavior. Early systemic immune activation by LPS was confirmed by increased spleen weight and serum cytokines. In brain tissue, gene expression analysis revealed a time course of inflammatory immune activation in TBI or LPS-treated mice (e.g., IL-1β, IL-6, CCL2, TNFα), which was exacerbated in TBI+LPS mice. This group also presented with fecal microbiome dysbiosis at 24 h post-LPS, with reduced bacterial diversity and changes in the relative abundance of key bacterial genera associated with sub-acute neurobehavioral and immune changes. Chronically, TBI induced hyperactivity and cognitive deficits, brain atrophy, and increased seizure susceptibility, similarly in vehicle and LPS-treated groups. Together, findings suggest that an immune challenge with LPS early after TBI, akin to a hospital-acquired infection, alters the acute neuroinflammatory response to injury, but has no lasting effects. Future studies could consider more clinically-relevant models of infection to build upon these findings.

## INTRODUCTION

Hospital-acquired infections represent a considerable challenge in the clinical management of patients following a severe traumatic brain injury (TBI).^1^ Mounting evidence suggests that the occurrence of such infections in TBI patients constitutes a dual-hit insult, whereby the infection and subsequent immune response amplifies the complexity of the injury response. This often results in extended hospital stays, additional complications including an increased risk of post-traumatic epilepsy (PTE)^2^, and poorer functional outcomes.^3-5^ Understanding the nuanced interplay between infections and TBI is paramount in elucidating the underlying mechanisms driving these deleterious synergies, and uncovering potential therapeutic targets to intervene and promote improved outcomes.^1^

Central to this scenario is the understanding that interactions between infections and TBI are predominantly mediated through inflammatory and immune responses. These responses, orchestrated by a multitude of cellular and molecular players including cytokines and chemokines, intricately shape the pathological trajectory that follows a TBI. Such immunological changes are thought to predispose patients to an increased risk of infection, while hospitalization and the need for often-invasive procedure to clinically manage a TBI also represent a distinct set of risk factors for infection.^1, 6^

Animal models serve as an invaluable tools in translational research, offering a controlled environment to dissect the complex interplay between infections and TBI. Using various approaches including lipopolysaccharide (LPS), poly I:C, or live bacteria models to mimic an infection, these models provide invaluable insights into the mechanistic underpinnings of how infections and a brain injury interact.^7^

In this study, we sought to elucidate the effects of a combined insult—experimental TBI and LPS to mimic a hospital-acquired infection post TBI—on a range of outcomes across an extended follow-up time course, from acute (hours) to chronic (months post-injury). In particular, we focused on the impact of TBI and LPS on neuroinflammation, the brain-gut axis, neurobehavioral manifestations, and the development of PTE, as important mechanisms and consequences of injury. In a previous study, we established a similar model paradigm in juvenile (post-natal day 21) male mice, and found that LPS induced a transient immune response but did not exacerbate chronic outcomes.^8, 9^ In the current study, we maintained a similar experimental timeline but used young adult mice (male and female), and a different source of LPS (*Klebsiella pneumoniae* instead of *Escherichia coli*), in line with another recent study from our group in which live *K. pneumoniae* were administered intratracheally as the infection model.^10^

## METHODS

### Animals and Ethics

All experimental procedures adhered to the guidelines stipulated in the Australian Code for the Care and Use of Laboratory Animals, as set forth by the National Health and Medical Research Council of Australia (NHMRC). Ethical approval for the experiments was obtained from both the Alfred Research Alliance Animal Ethics Committee and the Animal Care and Use Review Office of the Office of Research Protections, US Department of Defense. Male and female C57Bl/6J mice were sourced from the Walter and Eliza Hall Institute of Medical Research in Melbourne, Australia. Upon arrival, the mice were acclimatized for a period of one week prior to commencement of experimental procedures. All animal experimentation took place within dedicated facilities located in the Department of Neuroscience at the Central Clinical School, Monash University. Mice were group-housed in Optimice® individually-ventilated cages, with 2 to 6 mice per cage, under standardized conditions. The housing environment maintained a 12-hour light/dark cycle, with *ad libitum* access to both food and water throughout the duration of the study.

### TBI Model

Moderate to severe traumatic brain injury (TBI) was induced in male and female mice aged 10-11 weeks utilizing the Controlled Cortical Impact (CCI) model, following established protocols as previously described.^8-10^ Briefly, anesthesia was initiated with 4% isoflurane in oxygen, sustained via a nose cone at 1.5%. Preemptive analgesia was administered to all animals in the form of buprenorphine (0.05 mg/kg in saline, subcutaneously administered in the flank) and bupivacaine (1 mg/kg in saline, subcutaneously administered in the scalp) before the onset of surgery. Additionally, 0.5 mL of 0.9% saline was administered at the conclusion of the procedure to ensure adequate hydration.

Under stabilization within a stereotaxic frame, a midline incision was performed to expose the skull, followed by the creation of a 3 mm craniotomy in the left parietal bone to expose the intact dura mater. Employing an electronic controlled cortical impactor device (eCCI-6.3; Custom Design and Fabrication Inc., Sandston, VA), an impact was delivered at a velocity of 4.5 m/s, penetrating to a depth of 1.7 mm, and with a duration of 150 ms. Sham-operated animals underwent an identical surgical procedure without the brain impact. After the TBI or sham procedure, the incision site was sutured, topical antiseptic solution was applied, and mice were allowed to convalesce in individual cages atop a warming pad before being returned to their home cage.

### Lipopolysaccharide (LPS) Administration

On the fourth day after TBI or sham surgery (Figure 1a), animals were administered a 1 mg/kg intraperitoneal injection of LPS derived from *K. pneumoniae* (L4268, Sigma, Australia), prepared as a 0.2 mg/mL solution, or an equivalent volume of saline, to provoke an immune challenge. The chosen dose was determined based on established literature, which has shown that a 1 mg/kg intraperitoneal dose of LPS induces a transient immune response without resulting in mortality.^10^ Animals were subjected to close observation for indications of illness, such as manifestations of distress, lethargy, alterations in coat condition, or weight loss, utilizing a bespoke monitoring protocol following the injection.

**Figure 1:**
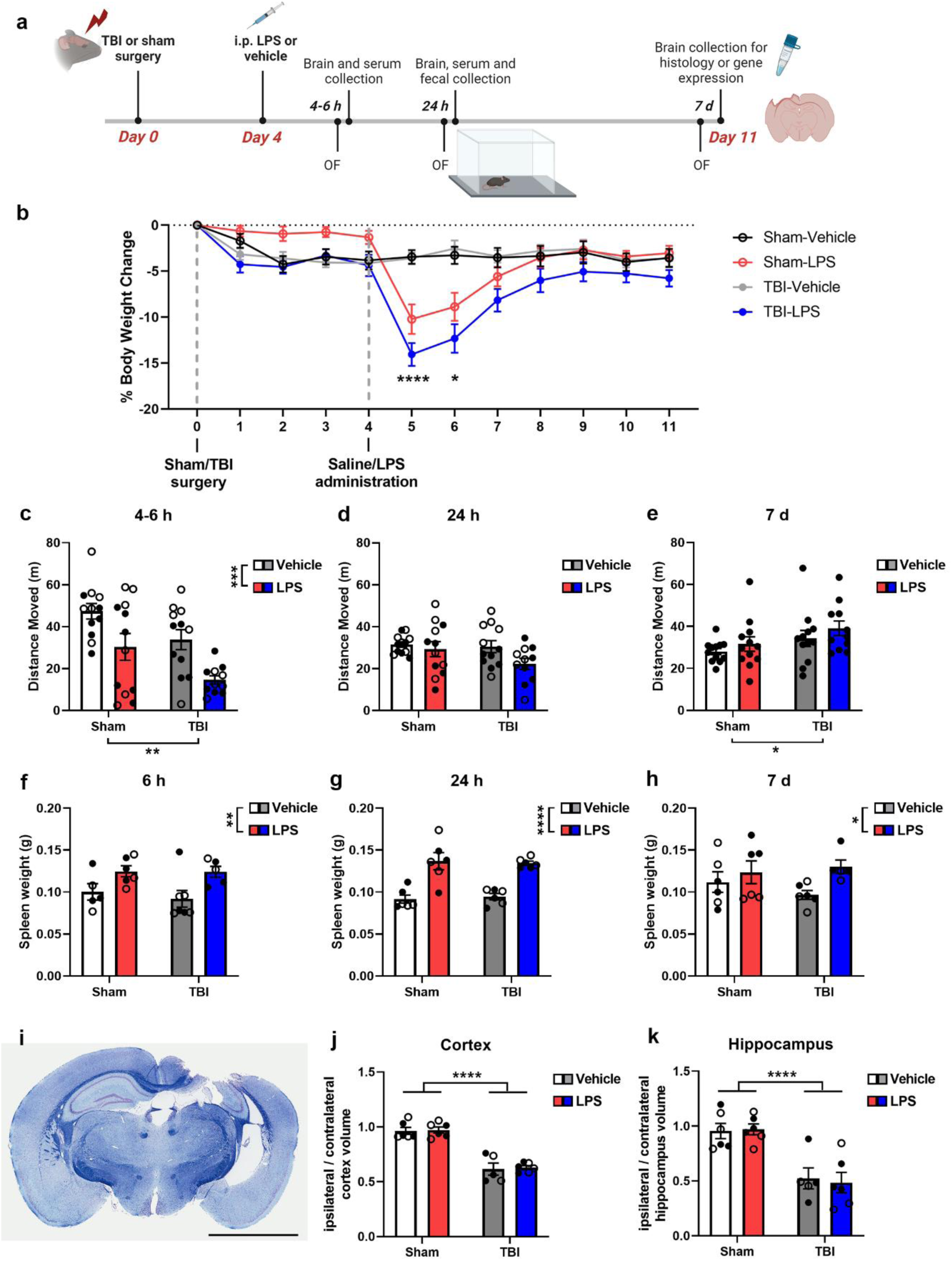
LPS post-TBI altered body and spleen weights and locomotor activity up to 7 days post-LPS, but did not alter the extent of tissue damage. LPS (1 mg/kg i.p.) was administered at 4 days post-TBI (a). Body weights were evaluated across the time course (b). For a cohort of mice that were euthanized at 7 d post-LPS, general activity was assessed in the open field test at 4-6 h (c), 24 h (d) and 7 d (e) post-LPS. Subsets of mice were euthanized at 6 h (f), 24 h (g) and 7 d (h) post-LPS. At 7 d post-LPS (i.e. 11 days post-TBI), dorsal cortex and hippocampal volumes were measured as an indicator of brain tissue loss. Scale bar = 3 mm. Open circles = females; closed circles = males. 2 or 3-way ANOVAs with post-hoc as appropriate; main effects are annotated under the x-axis or alongside the graph key, while post-hoc comparisons (where appropriate) are annotated within graphs. Timeline in panel (a) created in BioRender, Semple, B. (2022) BioRender.com/n47d527.

### Behavior Testing

At designated intervals of 6 hours, 24 hours, and 7 days post-administration of LPS or vehicle, all mice underwent assessment in an Open Field (OF) arena to gauge locomotor activity and anxiety-like behavior.^8, 10^ A 10-minute period of unrestricted exploration within the OF arena was recorded using an overhead camera and subsequently analyzed utilizing TopScan Lite v2.0.0 software (CleverSys Inc., Reston, VA, USA). Parameters measured included the total distance traversed and the proportion of time spent within a predefined central zone constituting 70% of the total arena area.

Upon reaching 14-16 weeks post-injury, a comprehensive battery of neurobehavioral tests was conducted to evaluate the enduring repercussions of TBI and LPS exposure. Following the OF assessment, anxiety-like behavior was further assessed through the employment of the Elevated Plus Maze (EPM) during a 10-minute session. The time spent in the open arms versus the closed arms of the maze was recorded using TopScan software.^8, 10^

Gross sensorimotor function was evaluated via the accelerating rotarod test administered across three consecutive days. Each day comprised three successive trials with a 30-minute inter-trial interval. The rotarod apparatus incrementally accelerated from 4 to 40 rotations per minute over a 5-minute period, with the latency to fall from the rotarod averaged per mouse per day.^8, 10^

Social approach and social novelty preferences were examined utilizing the three-chamber test, conducted across three consecutive 10-minute sessions within custom-built Perspex apparatus. This paradigm involved a habituation stage, followed by the introduction of a same-sex stimulus mouse into one outer chamber, and subsequently, the introduction of a novel same-sex stimulus mouse into the opposite outer chamber. TopScan software tracked the time spent by the experimental mouse in each outer chamber, with preferences indicative of social interest and recognition or memory.^8, 10^

Subsequently, the Morris Water Maze (MWM) test was employed to assess spatial learning and memory. A 120 cm diameter grey fiberglass tank filled with semi-opaque white-colored water served as the testing apparatus. Initial sessions, termed "visible platform" sessions, involved two daily sessions consisting of three 60-second trials each, with the platform elevated just above the water surface and marked with a flag. Following this, "hidden platform" sessions were conducted over days 2 to 5, comprising six trials per day without the flag. Mice relied on external 3D cues positioned around the pool to locate the submerged platform. TopScan software recorded latency to reach the platform and distance traveled. On day 8, a single "probe trial" was conducted with the platform removed, during which time spent in the target quadrant indicated spatial memory retention.^8, 10^

Lastly, the sucrose preference test was utilized to assess potential depressive-like anhedonia. Over a 5-day period, mice were presented with two drinking bottles containing either filtered water or a 1% sucrose solution. Bottle positions were alternated after 48 hours, and volumes consumed were measured on the first and last day of the test, with the sucrose preference ratio calculated as the volume of sucrose solution consumed divided by the total liquid consumed.^8, 10^

Behavioral data was captured from Sham-Vehicle (n=17), Sham-LPS (n=15), TBI-Vehicle (n=21), and TBI-LPS (n=23) mice. One mouse was excluded from the EPM and three-chamber tests due to tracking errors, and 5 mice were excluded from the sucrose preference test due to leakage of the water bottles.

### Video Electroencephalography (EEG)

To evaluate chronic seizure activity, electroencephalogram (EEG) electrodes were surgically implanted at 18-19 weeks post-injury, as described previously.^8, 11^ Under isoflurane anesthesia, epidural recording electrodes (E363/20/2.4/SPC ELEC W/SCREW SS, Plastics One Inc., USA) were positioned: one distal to the craniotomy, one contralateral to the craniotomy (2.5 mm right of the midline, -2.5 mm relative to Bregma), and two electrodes placed left or right over the cerebellum to serve as ground and reference, respectively. An additional anchor screw (00-96 x 3/32, Plastics One Inc., USA) was affixed over the left frontal region to reinforce the head cap, and all screws were securely attached to the skull using SuperGlue (Bostik, Australia). Subsequently, the electrodes were inserted into a pedestal head cap (MS363, Plastics One Inc. USA) and affixed with dental acrylate. The skin was sutured around the head cap, and animals received subcutaneous pain relief (buprenorphine 0.05 mg/kg and bupivacaine 1 mg/kg) and saline for hydration. Post-electrode insertion, mice were individually housed for the duration of the experiment.

Between weeks 20-24 post-TBI, continuous video-EEG recordings were captured using a Grael EEG amplifier device (Compumedics, Australia). EEG data was acquired unfiltered and digitized at sampling rate of 512 Hz with Profusion v4.0 (Compumedics, Australia). Acquisition employed high-pass (1 Hz) and low-pass (70 Hz) filters, Each animal underwent 10-16 days of video-EEG recording. Spontaneous seizures were assessed over a period of at least 10 days of recording per animal. Seizures were identified using the automated rodent seizure detection software ASSYST,^12^ and characterized by alterations in EEG patterns lasting >10 s, amplitude exceeding >3 times baseline, presenting as repetitive, rhythmic discharges with evolving amplitude.^11, 13^ An experienced investigator corroborated suspected seizures.

Immediately before tissue collection at 24 weeks post TBI, animals underwent video-EEG recording and received a single intraperitoneal dose of 40 mg/kg pentylenetetrazol (PTZ) (P6500, Sigma, Australia) to evaluate susceptibility to evoked seizures. Behavioral responses were monitored over a 15-minute period following PTZ administration and scored using a modified 7-point scale, where 0 indicates no response/normal activity, and 7 indicates status epilepticus culminating in mortality. From video-EEG recordings, the time to epileptiform discharge, time to generalized seizure onset, number of epileptiform spikes, and seizure duration were documented.^8, 13, 14^

Video-EEG data was captured from Sham-Vehicle (n=13), Sham-LPS (n=13), TBI-Vehicle (n=19), and TBI-LPS (n=21) mice. This excluded 2 mice that died after EEG electrode implantation, and one mouse that was euthanized at 21 weeks post-injury dye to rapid weight loss caused by undiagnosed malocclusion. Additional mice were excluded from EEG recordings due to their head caps detaching either before or during the recording period.

### Tissue Collection

Mice were euthanized using an intraperitoneal injection of 160 mg/kg sodium pentobarbitone (Lethabarb®, Virbac, Australia), followed by a cardiac blood collection. The blood samples were left to clot at room temperature for a minimum of 30 minutes before undergoing centrifugation to isolate serum for subsequent cytokine analysis. Following euthanasia, the spleen was promptly excised and weighed. Subsequently, mice were decapitated, and their brains were carefully removed and dissected. Tissue specimens were rapidly frozen in liquid nitrogen for subsequent gene expression analysis.

A subgroup of animals underwent euthanasia for brain sample collection for histological analyses, either at 11 days post-TBI or 24 weeks post-TBI (n=5-6/group/time point). These animals underwent transcardial perfusion with sterile saline, followed by 4% paraformaldehyde (PFA) at a rate of 2 mL/min. The extracted brains were subsequently post-fixed overnight in 4% PFA and then transferred to 70% ethanol before being sent to the Monash Histology Platform for paraffin processing and embedding (Monash University, Clayton, Australia). Coronal brain sections, with a thickness of 7 μm, were obtained, spanning approximately -0.7 mm to -3.4 mm Bregma.

### Microbiome Analysis

To investigate the impact of traumatic brain injury (TBI) and/or lipopolysaccharide (LPS) exposure on the fecal microbiome, fecal samples were obtained at 24 hours post-LPS/vehicle administration (equivalent to 5 days post-TBI/sham) for shallow shotgun whole-genome sequencing conducted by Transnetyx (Cordova, TN, USA). Briefly, individual mouse fecal samples (n=6 per group) were placed in uniquely barcoded sample collection tubes containing DNA stabilization buffer provided by Transnetyx, ensuring reproducibility, stability, and traceability. Subsequently, samples were shipped for DNA extraction, library preparation, and sequencing by Transnetyx.

DNA extraction was conducted utilizing the Qiagen DNeasy 96 PowerSoil Pro QIAcube HT extraction kit and protocol, employing an optimized and fully automated process to yield high-quality, inhibitor-free genomic DNA capturing the authentic microbial diversity of fecal specimens. Following extraction and quality control (QC), genomic DNA underwent conversion into sequencing libraries utilizing the KAPA HyperPlus library preparation protocol, a methodology designed to minimize bias. Unique dual-indexed (UDI) adapters were employed to prevent misassignment of reads and/or organisms. Upon completion of QC, libraries were subjected to shotgun sequencing on the Illumina NovaSeq platform, generating paired-end reads of 2x150 bp to a depth of 2 million read pairs.

Raw sequences were processed using the Sunbeam metagenomics pipeline.^15^ Shotgun index adapter sequences were removed using Trimmomatic,^16^ low-quality reads were removed, low-complexity reads filtered out using the pipeline’s Komplexity tool, and host reads identified and removed by mapping to the *Mus musculus* GRCm38 reference genome (NCBI BioProject: PRJNA20689). Bacterial sequences were assigned a taxonomy using the Kraken 2 tool^17^ with the complete RefSeq bacterial genomes database. Samples with fewer than 10,000 reads were excluded from the dataset and taxa present in fewer than 25% of samples or lacking assignment at the Phylum level were filtered out. Shannon index alpha diversity was determined using the estimate_richness function of the phyloseq (version 1.36.0) package.^18^ Read counts were normalized using Cumulative Sum Scaling (CSS) using the calcNormFactors function from metagenomeSeq (version 1.34.0) R package followed by log transformation (logCSS normalized).^19^

### Cytokine Multiplex

Cytokine and chemokine concentration was quantified in Bronchoalveolar-lavage fluid (BALF) and serum samples using a ProcartaPlex^TM^ Mouse and Rat Mix & Match Panel (CB230) custom 12-plex kit (Thermo Fisher Scientific, Lot: 326924-000), for detection of CCL2, CXCL1, CRP, G-CSF, GM-CSF, ICAM-1, IFNγ, IL-1α, IL-1β, IL-6, IL-10, and TNFα. The assay was performed as per manufacturer’s specifications on a MSD SQ120MM (VB204) instrument, with curves analyzed by Belysa 1.1.0 with 5PL curve fit optimization. Samples were assayed in duplicate by Crux Biolabs (Bayswater, VIC, Australia).

### Brain Gene Expression

RNA was extracted from brain samples (injured cortex and hippocampus) at 6 h, 24 h and 7 d post-infection, using the RNeasy® Mini Kit (Qiagen, Hilden, Germany). RNA purity was determined using a QIAxpert spectrophotometer then converted to cDNA via a QuantiTect® Reverse Transcription Kit (Qiagen). High-throughput qPCR analysis was carried out by the Monash Health Translation Precinct (MHTP) Medical Genomics Facility, using the Fluidigm BioMark HD™ system (Standard BioTools Inc., San Francisco, CA, USA) in a Fluidigm 192.24 Dynamic Array™ Integrated Fluidic Circuit for Gene Expression. A 24-gene panel were analyzed, including the housekeeping genes (HKGs) *Ywhaz* (Mm01722325_m1), *Ppia* (Mm02342430_g1) and *Hprt* (Mm03024075_m1),^20^ as well as the following cytokines, chemokines, immune cell markers, and immune-regulatory genes: *Il1β* (Mm00434228_m1), *Il1α* (Mm00439620_m1), *Tnfα* (Mm99999068_m1), *Il6* (Mm00446190_m1), *Il10* (Mm01288386_m1), *Ccl2* (Mm00441242_m1), *Ccr2* (Mm00438270_m1), *Cxcl2* (Mm00436450_m1) and *Cxcr2* (Mm99999117_s1), *Cd45* (Mm01293577_m1) and *Cd86* (Mm00444543_m1), *Gfap* (Mm01253033_m1), *Aldh1l1* (Mm03048957_m1), *Arg1* (Mm00475988_m1), *Tmem119* (Mm00525305_m1), *Trem2* (Mm04209424_g1), *Nos2* (Mm00440485_m1), *Tgf1* (Mm01227699_m1), *Mapk1* (Mm00442479_m1), *Nfkb1* (Mm00476361_m1), and *Hmox1* (Mm00516005_m1). Relative gene expression was calculated using the 2^-ρρCT^ method then normalized to the average expression of the three stable reference genes.^10, 20^

### Immunofluorescence

Brain sections were stained using several markers to evaluate cellular immune responses: glial fibrillary acidic protein (GFAP) for astrocytes, myeloperoxidase (MPO) for neutrophils, CD68 for macrophages, and IBA-1 for microglia/macrophages. The antigen retrieval process involved incubating the sections with DAKO Target Antigen Retrieval Solution (diluted 1:10; Agilent Technologies, USA) in a water bath at 90 °C for 45 minutes. Afterward, the sections were blocked with 10% normal donkey serum in 0.1% Triton-X100 in PBS. Primary antibodies were then applied in a 5% normal donkey serum solution: rabbit polyclonal GFAP (1:1000; Thermo Fisher Scientific), goat polyclonal MPO (1:250; R&D Systems), rabbit polyclonal IBA-1 (1:1000; WAKO), and rat anti-mouse CD68 (1:1000; Biorad). Secondary antibodies used were donkey anti-rabbit AF 594, donkey anti-goat AF 488, or donkey anti-rat AF 488 (1:250; Thermo Fisher Scientific), and counterstaining was done with Hoechst dye (1:1000; Thermo Fisher Scientific). Autofluorescence was quenched by immersing the slides in 0.1% filtered Sudan Black in 70% ethanol, followed by mounting with DAKO fluorescent mounting media. Images were captured using a Nikon Eclipse Ti-E inverted microscope with an Andor Zyla sCMOS camera and Nikon NIS-Elements v.5.30.06 software at 20x magnification. Stitched images of either the entire lung section or the ipsilateral dorsal hemisphere were obtained and analyzed in FIJI ImageJ for manual cell counts (e.g., MPO cells in the injured brain) or fluorescence percentage area (for GFAP and IBA-1).^10^ A blinded investigator performed these analyses within specified regions of interest (ROIs), averaging the results across 2 sections per brain per marker.

### Histology

Brain tissue underwent Luxol Fast Blue (LFB) and Cresyl Violet (CV) staining following a modified established protocol.^10^ Initially, the tissue was subjected to dewaxing in xylene, rehydration in 95% ethanol, and immersion in LFB solution at 40 °C for a duration of 6 hours. Subsequent steps included washing the tissue in 70% ethanol and distilled water for 20 seconds, differentiation in a 0.01% lithium carbonate solution, followed by immersion in 70% ethanol for 20 seconds and rinsing with distilled water. The tissue was then stained with CV acetate (0.5% w/v solution in 0.5% w/v acetic acid) for 10-20 minutes, immersed in absolute ethanol thrice for 1 minute each, briefly exposed to xylene twice for 2 minutes each, and finally mounted with DPX Mountant (Sigma-Aldrich, St. Louis, MO, USA).

Staining with LFB and CV was carried out on 7 equidistant sections, spaced 280 µm apart, covering approximately -0.7 to -3.5 mm Bregma. Tissue atrophy was quantified utilizing previously established methods. Digital imaging was conducted utilizing a Leica Aperio AT Turbo Brightfield slide scanner located at the Monash Histology Platform. The acquired images were exported to FIJI/ImageJ software for analysis, employing the unbiased Cavalieri method to estimate the volume of the intact dorsal cortex and dorsal hippocampus in each hemisphere (ipsilateral and contralateral to the injury).^21, 22^ Each grid point represented an area of 100 µm², with slides sampled at intervals of four. Measurements were confined to the dorsal aspect of each region of interest. Group means were represented as the ratio of ipsilateral to contralateral volumes.

### Statistical Analysis

Statistical analyses were conducted using GraphPad Prism v.9.4.1 (GraphPad Software Inc., San Diego, CA, USA), with a significance level set at p<0.05. Two and three-way analyses of variance (ANOVA) were employed, complemented by Tukey’s post hoc test where applicable. Data featuring three independent variables (time, injury, and LPS) underwent analysis by 3-way ANOVA. In most instances, male and female mice were pooled within each group, with open circle data points specifically representing female animals. Potential sex differences were evaluated via 3-way ANOVA, considering the factors of sex, injury, and LPS, and reported solely if significant sex differences were observed. Microbiome data were analyzed using R (version 4.3.0). All results are presented as mean ± standard error of the mean (SEM). Differential abundance testing for bacterial taxa was achieved using linear modelling via a custom wrapper function around the limma R package (version 3.48.3)^23^ as detailed here: https://github.com/mucosal-immunology-lab/microbiome-analysis/wiki/Limma-DA. Benjamini-Hochberg multiple testing correction was used with a q-value significance threshold of 0.05. A fixed random-number seed value of 2 was set to ensure maximum reproducibility of tools requiring random pseudo-numbers.

## RESULTS

### LPS Administration Following TBI Led to Reduced Body Weights and Locomotor Activity

In the first experiments, male and female young adult mice underwent TBI or sham surgery on day 0, followed by LPS or vehicle at day 4, and analysis of tissue collected across the first week thereafter to examine potential interactions between TBI and LPS (Figure 1a). Changes in body weight over time were quantified as a measure of general health (Figure 1b). Following TBI/sham surgery, while most groups appeared to drop up to 5% of their baseline body weight, this did not reach statistical significance (3-way ANOVA main effect of TBI, F_1, 43_=3.67 p=0.0622). LPS did affect body weights (main effect of LPS, F_1, 43_=5.60 p=0.0226), and a significant interaction between time and LPS was observed (F_1, 43_=29.80 p<0.0001). Groups that received LPS on day 4 exhibited significant weight loss on day 5 and 6 (p<0.0001 and p<0.05) compared to groups that received vehicle, but regained weight from day 7 onwards.

General locomotor and exploratory activity of mice was observed in the OF test at 6 h, 24 h and 7 d following LPS or vehicle administration (Figure 1c-e). At 6 h, both TBI and LPS independently reduced the distance moved in the OF (2-way ANOVA main effect of TBI, F_1,43_=10.21 p=0.0026; main effect of LPS, F_1,43_=15.58 p=0.0003), which resolved by 24 h post-LPS. By 7 d post-LPS, a modest effect of injury was observed (F_1,43_=4.65 p=0.0367), whereby TBI mice moved a greater distance compared to sham mice. No additive effect of TBI and LPS was observed.

Fresh spleen weights were measured in a subset of mice at each time point, as an indirect indicator of immune activation (Figure 1f-h). LPS induced an increase in spleen weight at 6 h (2-way ANOVA F_1,19_=10.45 p=0.0044), 24 h (F_1,19_=49.34 p<0.0001) and 7 d post-LPS (F_1,19_=4.58, p=0.0455). However, TBI had no overt effect on spleen weights, nor were there any TBI x LPS interactions.

Finally, the extent of cortical and hippocampal damage was evaluated at 7 d post-LPS from histologically-stained coronal brain sections (Figure 1i). TBI resulted in considerable volume loss in the injured cortex compared to sham controls (Figure 1j, 2-way ANOVA F_1,19_ =105.4 p<0.0001), with no differences between vehicle and LPS treatment. Similarly, the ipsilateral hippocampal volume was reduced by TBI only (Figure 1k, 2-way ANOVA F_1,19_=34.46 p<0.0001).

### LPS Administration Triggered a Robust Systemic Cytokine Response

Serum was analyzed to evaluate the immune response to TBI and LPS administration at 6 h post-LPS/vehicle (on the fourth day post-TBI/sham) (Table 1). No effects were seen in sham or TBI only groups. In contrast, LPS administration increased IL-1β, IL-6, IL-10, CCL2, CXCL1, G-CSF, CRP and ICAM1 in both LPS-treated groups. No combinatory effects of TBI and LPS were observed (two-tailed unpaired t-tests comparing Sham-LPS vs. TBI-LPS; p>0.05). IFNγ, TNFα, IL-1α and GM-CSF were below detectable levels in the majority of samples, and thus were not analyzed.

**Table 1:**
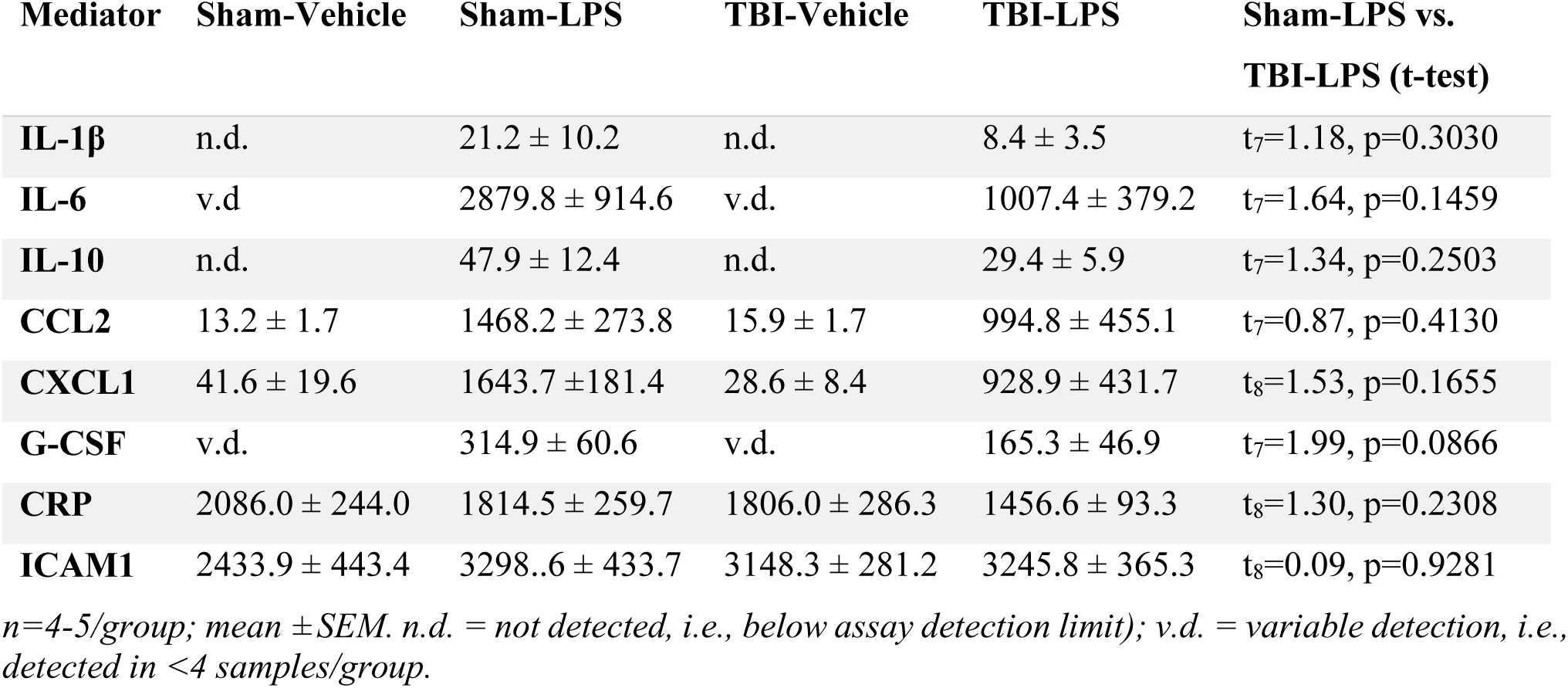
Levels of Soluble Mediators in Serum at 6 h Post-LPS (pg/mL)

| Mediator | Sham-Vehicle | Sham-LPS | TBI-Vehicle | TBI-LPS | Sham-LPS vs.<br>TBI-LPS (t-test) |
| --- | --- | --- | --- | --- | --- |
| **IL-1β** | n.d. | 21.2 ± 10.2 | n.d. | 8.4 ± 3.5 | t<sub>7</sub>=1.18, p=0.3030 |
| **IL-6** | v.d. | 2879.8 ± 914.6 | v.d. | 1007.4 ± 379.2 | t<sub>7</sub>=1.64, p=0.1459 |
| **IL-10** | n.d. | 47.9 ± 12.4 | n.d. | 29.4 ± 5.9 | t<sub>7</sub>=1.34, p=0.2503 |
| **CCL2** | 13.2 ± 1.7 | 1468.2 ± 273.8 | 15.9 ± 1.7 | 994.8 ± 455.1 | t<sub>7</sub>=0.87, p=0.4130 |
| **CXCL1** | 41.6 ± 19.6 | 1643.7 ± 181.4 | 28.6 ± 8.4 | 928.9 ± 431.7 | t<sub>8</sub>=1.53, p=0.1655 |
| **G-CSF** | v.d. | 314.9 ± 60.6 | v.d. | 165.3 ± 46.9 | t<sub>7</sub>=1.99, p=0.0866 |
| **CRP** | 2086.0 ± 244.0 | 1814.5 ± 259.7 | 1806.0 ± 286.3 | 1456.6 ± 93.3 | t<sub>8</sub>=1.30, p=0.2308 |
| **ICAM1** | 2433.9 ± 443.4 | 3298.6 ± 433.7 | 3148.3 ± 281.2 | 3245.8 ± 365.3 | t<sub>8</sub>=0.09, p=0.9281 |

*n=4-5/group; mean ± SEM. n.d. = not detected, i.e., below assay detection limit); v.d. = variable detection, i.e., detected in <4 samples/group.*

A similar pattern of the above-mentioned soluble cytokines, chemokines and immune-related proteins was evident at 24 h post-LPS/vehicle (=5 days post-TBI/sham), with LPS appearing to stimulate protein levels but to a similar extent in TBI and Sham mice. Mediator levels were typically lower overall by this time point (data not shown).

### Independent and Additive Effects of TBI and LPS on Brain Inflammatory Gene Expression

Turning to the injured brain, we sought to understand whether the secondary immune challenge of LPS would alter the neuroinflammatory response to TBI. The expression of a panel of immune-related genes was evaluated by qPCR across the time course (6 h, 24 h and 7 d). Figure 2a illustrates gene expression patterns at 6 h post-LPS. TBI alone resulted in an increase in prototypical inflammatory/immune genes including *Gfap, CD86* and *Trem2.* LPS alone increased gene expression of cytokines *Il1α, Il6, Il1β* and *Tnfα*, as well as the chemokine *Cxcl2* and its receptor *Cxcr2*. Of particular interest, however, were the interactions between TBI and LPS, detected for inflammatory cell surface markers including *Cd45, Ccr2* and *Cxcr2,* cytokines/chemokines including *Il1β, IL1α, Tnfα, Il6* and *Ccl2*, and intracellular signaling molecules *Nfkb1, Nox2* and *Hmox1* (2-way ANOVAs, TBI x LPS interaction p<0.05). Expression patterns were similar at 24 h, albeit typically a magnitude lower, while by 7 d the effects of LPS were resolved.

**Figure 2:**
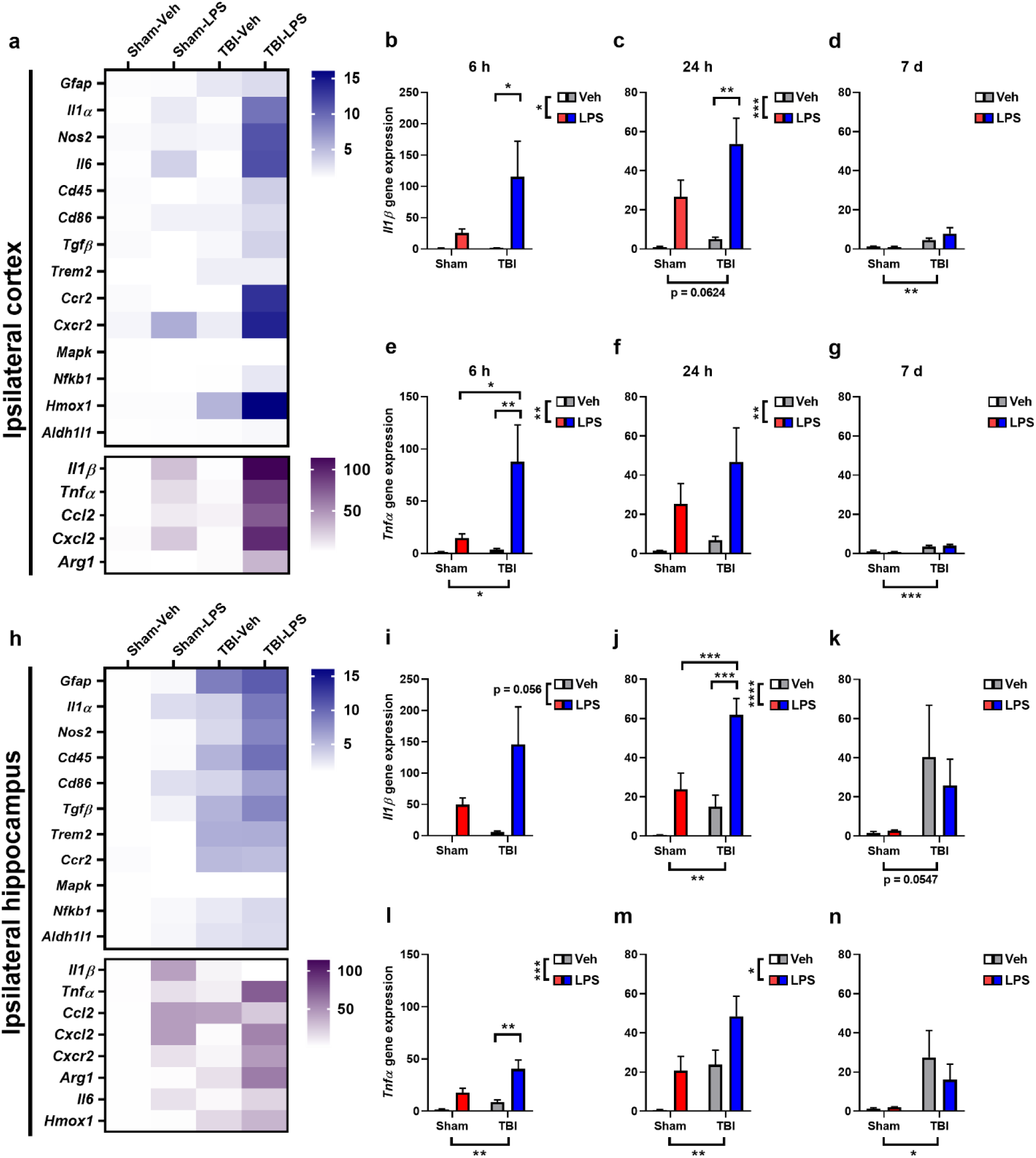
Gene expression analysis in the ipsilateral injured brain revealed an additive effect of TBI + LPS for several key inflammatory mediators. qPCR was performed at 6 h, 24 h and 7 d post-LPS. Gene expression in the ipsilateral cortex at 6 h is depicted by a heat map (a), at a range of 0-15 (blue, top) or 0-100 (below, purple) depending on the relative expression levels. Expression of Il1β is presented graphically at 6 h (b), 24 h (c) and 7 d post-LPS (d), to illustrate the time course; followed by Tnfα (e-g). In the lower panels, gene expression in the ipsilateral hippocampus at 6 h is depicted by a heat map (h), alongside Il1β and Tnfα at 6 h (i, l), 24 h (j, m) and 7 d (k, n). n=4-5/group, 2 or 3-way ANOVAs with post-hoc as appropriate; main effects are annotated under the x-axis or alongside the graph key, while post-hoc comparisons (where appropriate) are annotated within graphs.

This time course is exemplified by *Il1β* (Figure 2b-d) and *Tnfα* (Figure 2e-g). Gene expression of *Il1β* was increased by LPS at 6 h (2-way ANOVA, F_1,19_=7.75 p=0.0118), with trends towards a main effect of injury (F_1,19_=3.28 p=0.0859) and an LPS x TBI interaction (F_1,19_=3.24 p=0.0878). Sidak’s post-hoc comparisons revealed a significant difference between TBI-Vehicle and TBI-LPS groups (p<0.05). At 24 h, there was a robust main effect of LPS (F_1,20_=22.53 p=0.0001) and a trend towards a main effect of TBI (F_1,20_=3.90 p=0.0624); and again, higher levels of *Il1β* in TBI-LPS compared to TBI-Vehicle groups (post-hoc, p<0.01). By 7 d, the effect of LPS had resolved (F_1,19_=1.36 p=0.2573), but a subtle yet significant effect of TBI persisted (F_1,19_=13.12 p=0.0018). For *Tnfα*, main effects of TBI (F_1,19_=5.34 p=0.0329) and LPS (F_1,19_=9.0 p=0.0077), and a significant TBI x LPS interaction (F_1,19_=4.67 p=0.0445) were observed, with post-hoc comparisons highlighting higher gene expression in the TBI-LPS compared to TBI-Vehicle group (p<0.01). By 24 h, only a main effect of LPS was noted (F_1,20_=9.89 p=0.0051), while *Tnfα* expression at 7 d mirrored *Il1β* in that the LPS effect had resolved while a TBI-induced increase remained (F_1,19_=23.03 p=0.0002).

In the ipsilateral hippocampus, at 6 h post-LPS a robust pro-inflammatory response akin to that seen in the injured cortex was observed, with elevated expression of multiple cytokines, chemokines and immune cell markers (Figure 2h). For exemplar gene *Il1β*, a trend towards an LPS effect was seen at 6 h (Figure 2i; 2-way ANOVA F_1,14_=4.33, p=0.056). While the magnitude was lower by 24 h, at this time, effects of both LPS and TBI were noted (F_1,17_=23.61, p=0.0001 and F_1,17_=13.22, p=0.002, respectively). Post-hoc analyses indicating that *Il1β* expression in the injured hippocampus of TBI-LPS mice was higher than either TBI or LPS alone (Figure 2j; p=0.001 and p=0.0002, respectively). For *Tnfα*, main effects of both TBI and LPS were detected at 6 h (Figure 2l; F_1,15_=9.53, p=0.0075 and F_1,15_=24.43, p=0.0002, respectively) and 24 h (Figure 2m; F_1,17_=10.56, p=0.0047 and F_1,17_=8.17, p=0.0109, respectively). By 7 d, a modest effect of TBI persisted (F_1,16_=6.38, p=0.0225) while the effect of LPS had resolved (Figure 2n).

### Dysbiosis of the Fecal Microbiome in TBI+LPS Mice

Increasing evidence has uncovered close, bidirectional relationships between the brain, the immune system and microbiome, with several studies indicating an effect of TBI on the fecal microbiome.^24-27^ To explore whether the combined insult of experimental TBI and a subsequent LPS-mediated immune challenge altered the fecal microbiome, fecal samples were collected and analyzed for bacterial content.

At 24 h post-LPS/vehicle, alpha diversity was comparable in Sham-Vehicle, Sham-LPS and TBI-Vehicle groups; however, diversity was reduced in the fecal microbiome of TBI-LPS mice compared to TBI-Vehicle mice (Figure 3a; p=0.0043). When comparing microbiome composition at the phyla level, no overt differences in the relative abundance of different phyla were observed between Sham-Vehicle and Sham-LPS mice, or Sham-Vehicle and TBI-Vehicle mice (Figure 3b). However, the abundance of *Actinobacteria* and *Proteobacteria* appeared to be increased in TBI-LPS samples. Exploratory limma differential abundance analysis confirmed that the greatest effects were in regards to LPS treatment (compared to vehicle) in TBI animals (Figure 3e). Box plots to visualize relative abundance at the genera level revealed that the TBI-LPS group was characterized by an increase in *Curtobacterium* and *Proteus* alongside a concurrent decrease in *Prevotella* (Figure 3g).

**Figure 3:**
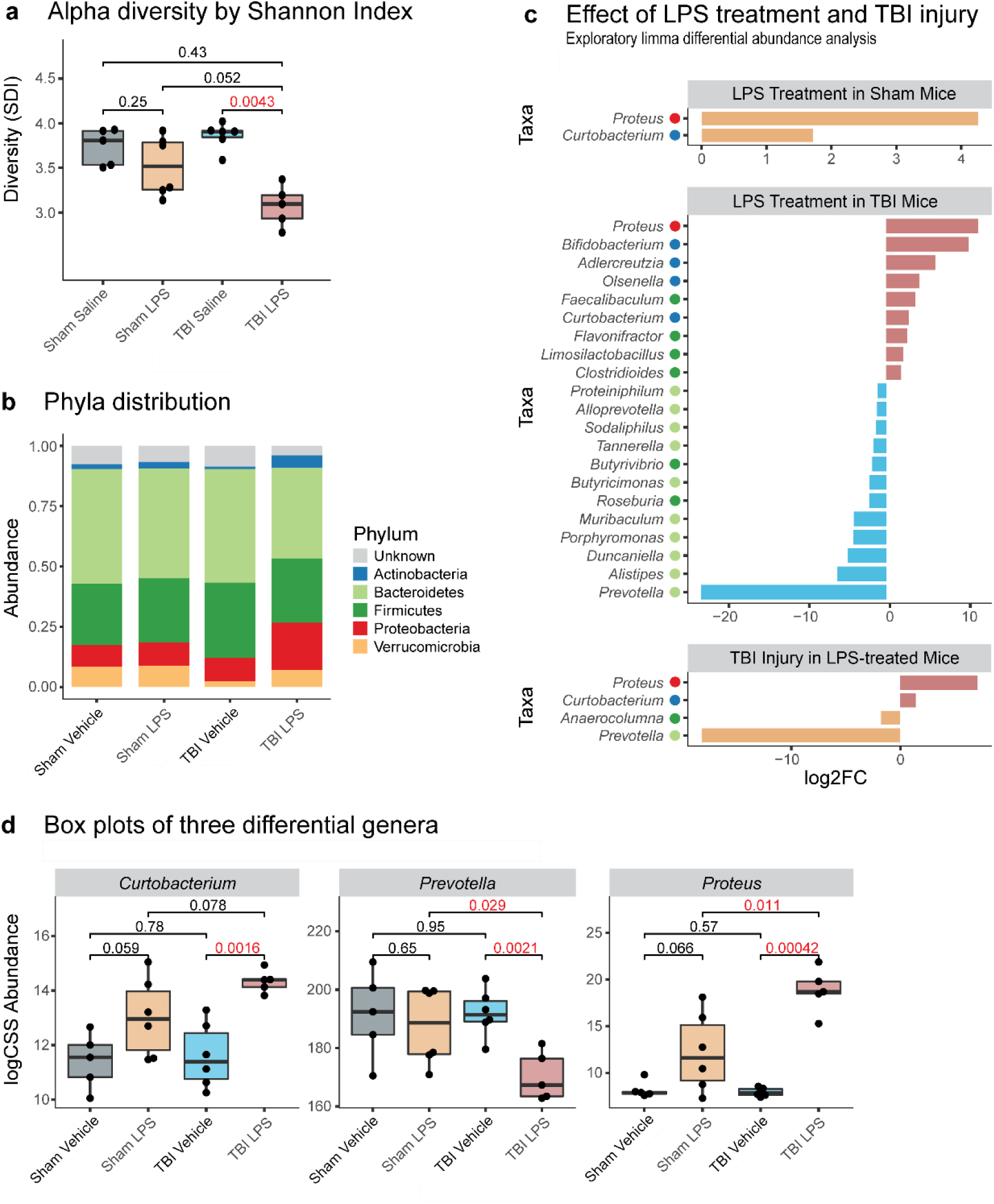
The combination of TBI + LPS alters the fecal microbiome diversity and composition. At 24 h post-LPS, alpha diversity was reduced in the fecal microbiome of TBI-LPS mice compared to TBI-Vehicle mice (a). No overt differences in relative abundance of different phyla were observed between Sham-Vehicle and Sham-LPS mice, or Sham-Vehicle and TBI-Vehicle mice (b). Exploratory limma differential abundance analysis confirmed the most pronounced effects were due to LPS treatment (compared to Vehicle) in TBI animals (c). Examples of significantly different genera *Curtobacterium*, *Prevotella* and *Proteus* are depicted as box plots in (d); analysis by Student’s t-test followed by Shapiro-Wilk test. n=5-6/group.

Delving into potential interactions between the brain and gut, linear modelling analyses were performed to ask whether any changes in the fecal microbiome were associated with behavior or immune responses. Significant associations were detected between changes in particular bacterial genera in the fecal microbiome and brain gene expression in the injured cortex and hippocampus. For example, an increase in *Proteus* was positively associated with higher levels of *CD45, Il1α, Il1β,* and several other inflammatory mediators in the injured brain. Correlations between fecal microbiome composition and behavioral measures were also observed; for example, higher abundance of *Prevotella* was associated with higher activity (greater distance travelled) in the open field task.

### Cellular Neuroinflammation Persists to at Least 11 Days after TBI and LPS

Finally, to test the hypothesis that experimental TBI followed by an LPS would exacerbate cellular neuroinflammation, leukocyte infiltration and glial reactivity were assessed in the injured brain (ipsilateral hemisphere) at 7 d post-LPS/Vehicle (11 days post-TBI/Sham). Iba1+ staining for microglia and macrophages was increased in the cortex and hippocampus, to a similar extent in both Vehicle and LPS-treated groups (Figure 4a-c; 2-way ANOVA main effect of TBI, F_1,20_=22.95, p=0.0001 and F_1,20_=27.12, p<0.0001, for cortex and hippocampus, respectively). Staining for CD68, presumably identifying infiltrating or activated macrophages, was less widespread and largely located in dense patches close to the lesion core (Figure 4d-f). A trend towards increased CD68 staining due to TBI was detected for the cortex (F_1,20_=4.21, p=0.0535), and a significant TBI effect was seen in the hippocampus (F_1,20_=10.04, p=0.0048). Next, the extent of GFAP staining was quantified as an indicator of astrocyte reactivity (Figure 4g-i). Again, only a main effect of TBI was observed at this time point in both the cortex (F_1,19_=34.41, p<0.0001) and hippocampus (F_1,19_=40.96, p<0.0001). Finally, MPO staining—presumed to represent infiltrating neutrophils based on the staining pattern and cellular morphology (polynucleated cells identified at high magnification)—was quantified from cell counts (Figure 4j-l). In the injured cortex, while no cells were detected in either Sham-Vehicle or Sham-LPS groups, main effects of both TBI (F_1,18_=19.46, p=0.0003) and LPS (F_1,18_=5.34, p=0.0329) were observed, as well as a significant TBI x LPS interaction (F_1,18_=5.34, p=0.0329). From post-hoc analysis, more MPO+ cells were counted in the TBI-LPS group compared to the TBI-Vehicle group (p<0.05). Similarly, in the injured hippocampus, main effects of TBI (F_1,18_=20.77, p=0.0002) and LPS (F_1,18_=0.0355) were observed, as well as a significant TBI x LPS interaction (F_1,18_=5.17, p=0.0355), driven again by an increased number of cells counted in the brains of TBI-LPS compared to TBI-Vehicle mice (p<0.05).

**Figure 4:**
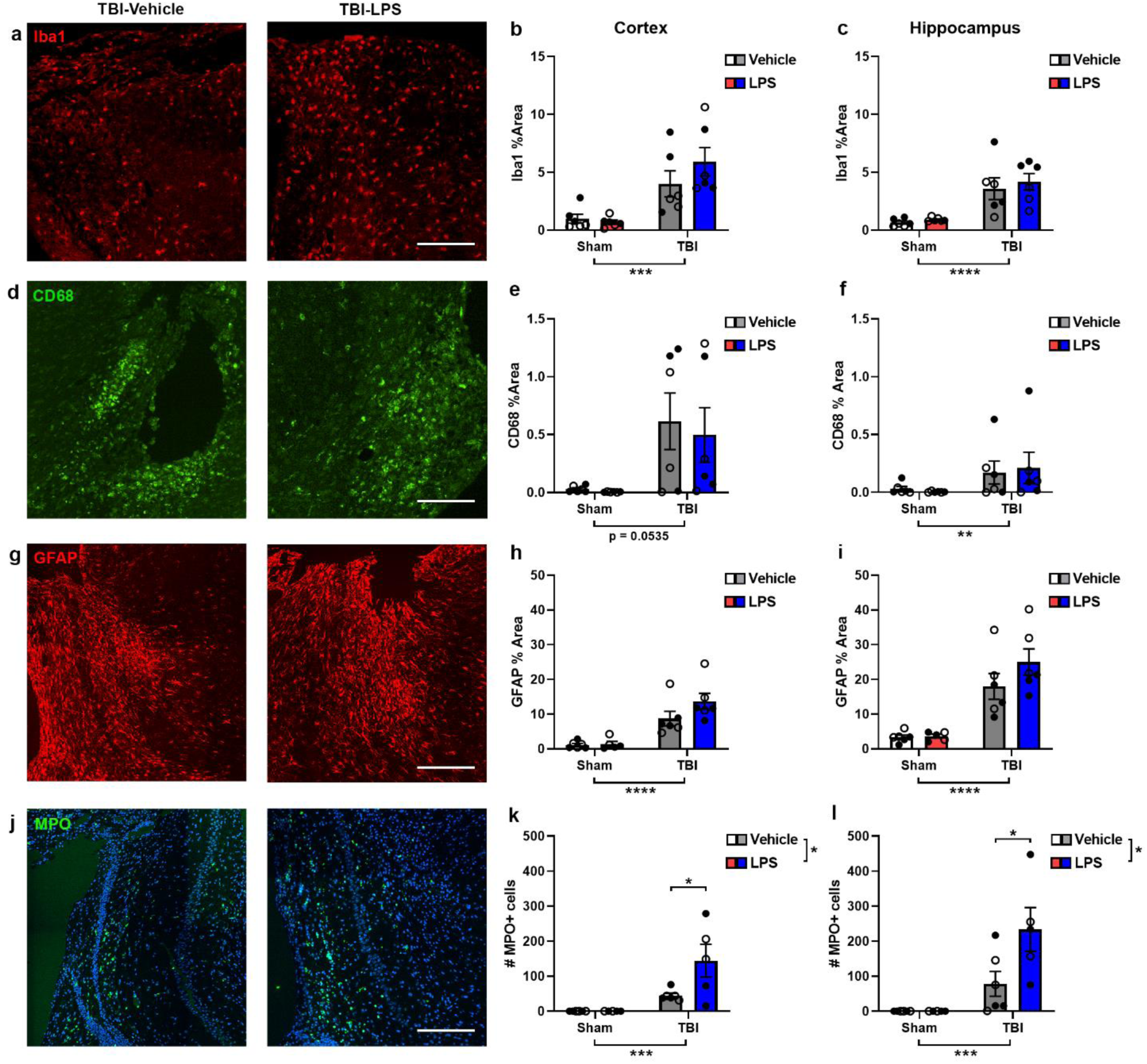
Leukocyte infiltration and glial reactivity in response to TBI and/or LPS. Immunofluorescence staining for Iba1 representing microglia (a-c), CD68 for macrophages (d-f), GFAP for astrocytes (g-i) and MPO for neutrophils (j-l) revealed robust immune cell activation and infiltration at 7 days post-LPS/Vehicle. Increased cell numbers or percentage area staining was predominantly driven by TBI, with an effect of LPS seen only for MPO+ cell numbers, which were higher in both the injured cortex and hippocampus of TBI-LPS compared to TBI-Vehicle mice. Open circles = females; closed circles = males. 2-way ANOVAs with post-hoc as appropriate; main effects depicted as brackets and bars between groups (outside graphs) and post-hoc results depicted within graphs. *p<0.05, **p<0.01, ***p<0.001, ****p<0.0001. n=5-6/group.

### Chronic Neurobehavioral Outcomes Revealed TBI Deficits Unaffected by LPS

To address the hypothesis that the combined insult of TBI and LPS would exacerbate chronic post-injury outcomes, an extended experimental timeline was established (Figure 5a). Mice received TBI or sham surgery on day 0, followed by LPS or Vehicle on day 4, as described above. Mice were then allowed to recover until weeks 14-16 post-injury, when they underwent a series of neurobehavioral tests. This was followed by surgical implantation of recording electrodes for a period of video-EEG monitoring to evaluate potential post-traumatic epilepsy.

**Figure 5:**
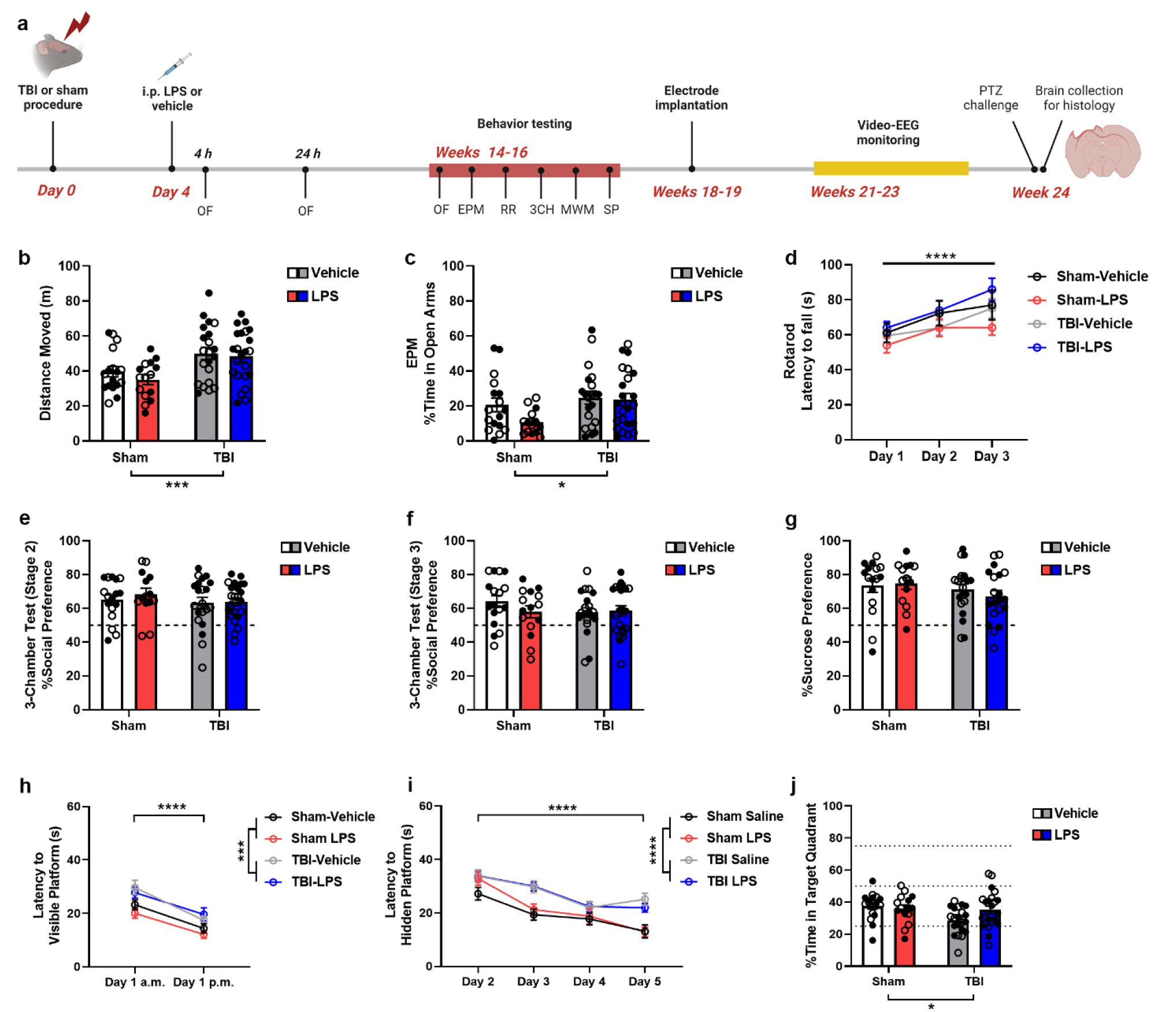
A chronic time course was established to evaluate long-term behavior, seizure, and neuropathological outcomes after TBI and LPS. The experimental timeline is depicted in (a). Created in BioRender, Semple, B. (2023) BioRender.com/g61z794. At 14-16 weeks post-injury, activity in the Open Field test (b) and Elevated Plus Maze (EPM; c) was altered by TBI only (Two-way ANOVA effect of injury, ***p = 0.0005 and *p=0.0473, respectively). In the accelerating rotarod (d), all groups improved in performance with subsequent trials (3-way ANOVA effect of time, ****p<0.0001). In the the three-chamber social approach task, following an initial habituation stage (not shown), no effects of either TBI or LPS were observed in stages 2 or 3 (e-f). Similarly, no change in sucrose preference were noted between groups (g). In the Morris Water Maze task, while all mice showed improvement on day 1 (the visible platform session) between the a.m. and p.m. sessions (h), TBI groups travelled a greater distance before reaching the platform (main effect of injury, **p=0.0094). Similarly, all groups showed decreased distance travelled before reaching the hidden platform on day 2-5 (i), with TBI animals travelled a longer distance (****p < 0.0001). Finally, a subsequent probe trial test (j) revealed that the time spent in the target quadrant was reduced in TBI groups (*p=0.0315). Dotted line in e, f, g and j indicates 50% preference (chance). Open circles = female; closed circles = male. n=15-17 (Sham groups) and 21-23 (TBI groups).

In the Open Field test (Figure 5b), TBI mice travelled a greater distance compared to Sham mice irrespective of LPS treatment (2-way ANOVA main effect of injury, F_1,72_=13.34, p=0.0005). In the EPM (Figure 5c), an effect of TBI was also observed, as an increase in the proportion of time spent by TBI groups in the open arms of the maze (F_1, 71_=5.53, p=0.0215). Together, these findings indicate that TBI results in a persistent hyperactive phenotype associated with a reduction in anxiety-like behavior. No effects of LPS were observed.

No effects of either TBI or LPS were observed in the accelerating rotarod test (Figure 5d), with all groups showing an improvement in sensorimotor performance (increased latency to fall) over subsequent trial days (3-way ANOVA F_2,144_=19.56, p<0.0001). Similarly, no effects of TBI or LPS were observed in the three-chamber social approach task, in either Stage 1 (habituation; not shown), Stage 2 (Figure 5e), or Stage 3 (Figure 5f), indicating that all groups showed intact sociability and social memory. Further, none of the experimental groups showed anhedonia, with all groups demonstrating a strong preference for sucrose over water in the sucrose preference test (Figure 5g).

The MWM was conducted over a one-week period to evaluate spatial learning and memory. In the initial visible platform trials (Figure 5h), all groups showed an improvement in performance (reduced latency to the platform) between the morning and afternoon sessions (3-way ANOVA F_1,72_=23.09, p<0.0001). A main effect of TBI was also observed (F_1,72_=7.13, p=0.0094), with TBI groups travelling a greater distance before reaching the platform. In the subsequent hidden platform sessions (Figure 5i), on days 2-5, we again saw improvement over time (F_2, 188_=41.85, p<0.0001) and a main effect of TBI (F_1,72_=26.56, p<0.0001). In addition, a significant TBI x time interaction (F_3,216_=3.14, p=0.0263) reflected the divergence of performance curves between Sham and TBI groups, whereby Sham groups showed continued improvement (reduced latency to find the platform) across the week, while TBI groups on days 4-5. Finally, a single probe trial was conducted to test for spatial memory retention (Figure 5j), a task that failed to differentiate the experimental groups which all showed a preference for the target quadrant (where the platform was previously located).

### TBI Led to Increased Susceptibility to Evoked Seizures and Chronic Neuropathology by 6 Months

Spontaneous and evoked seizures responses were evaluated at a chronic time point post-TBI/LPS to evaluate post-traumatic epilepsy. Spontaneous seizures were assessed over a period of at least 10 days of recording per animal (Figure 5a). No spontaneous seizure activity was observed in either Sham-Vehicle or Sham-LPS mice (Table 2). Two mice (10.53%) in the TBI-Vehicle group, and only one mouse in the TBI-LPS group (4.76%) showed at least one spontaneous seizure during the monitoring period (Figure 6a-b). The average seizure duration in TBI-Vehicle mice appeared to be higher than seizure duration in the TBI-LPS mouse (66.67 vs. 36 s), alongside a higher Racine Scale Score of 4 vs. 3. However, the low number of seizure events prevents statistical comparison.

**Figure 6:**
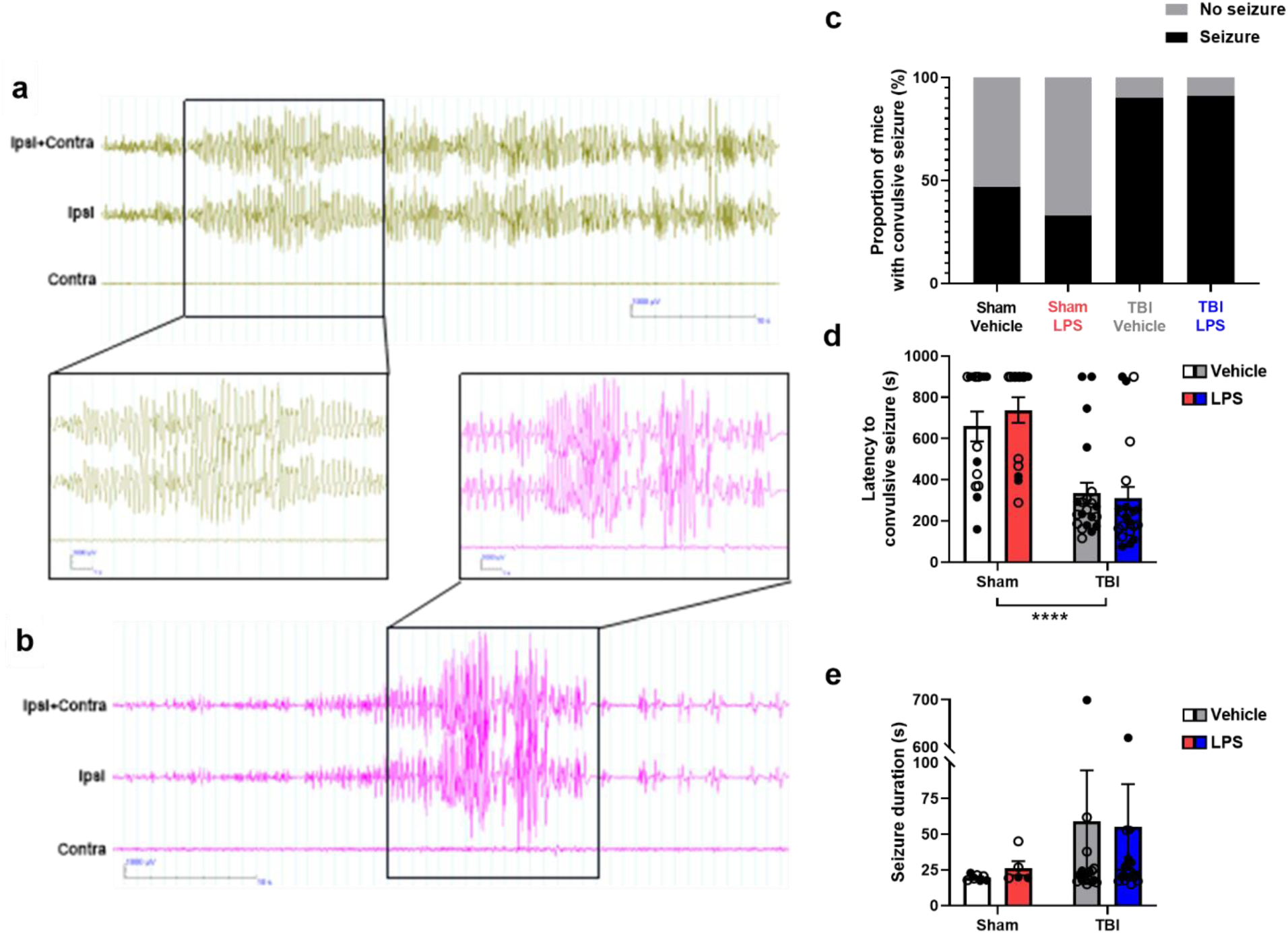
Chronic spontaneous seizures were rare, but TBI mice showed increased vulnerability to evoked seizures. Representative EEG recordings during spontaneous seizures from a TBI-Vehicle mouse (a) and TBI-LPS mouse (b); except illustrates a higher magnified view of part of the seizure. EEG activity viewed in Profusion EEG v.6, with high pass filter 1 Hz, low pass filter 30 Hz, and notch filter 50 Hz). Ipsi = electrode recording from the cortex ipsilateral to the site of TBI; Contra = contralateral cortex to the TBI. After recording for spontaneous seizure activity over 10 days, all mice received a low-dose PTZ challenge to evoke seizures. The proportion of mice that responded to PTZ with a convulsive seizure was higher in TBI compared to Sham groups (c). Latency to a convulsive seizure was reduced in TBI groups (d), while seizure duration was variable (e). Open circles = female; closed circles = male. ****p<0.0001; n=13/group (Sham) or 19-21/group (TBI).

**Table 2:**
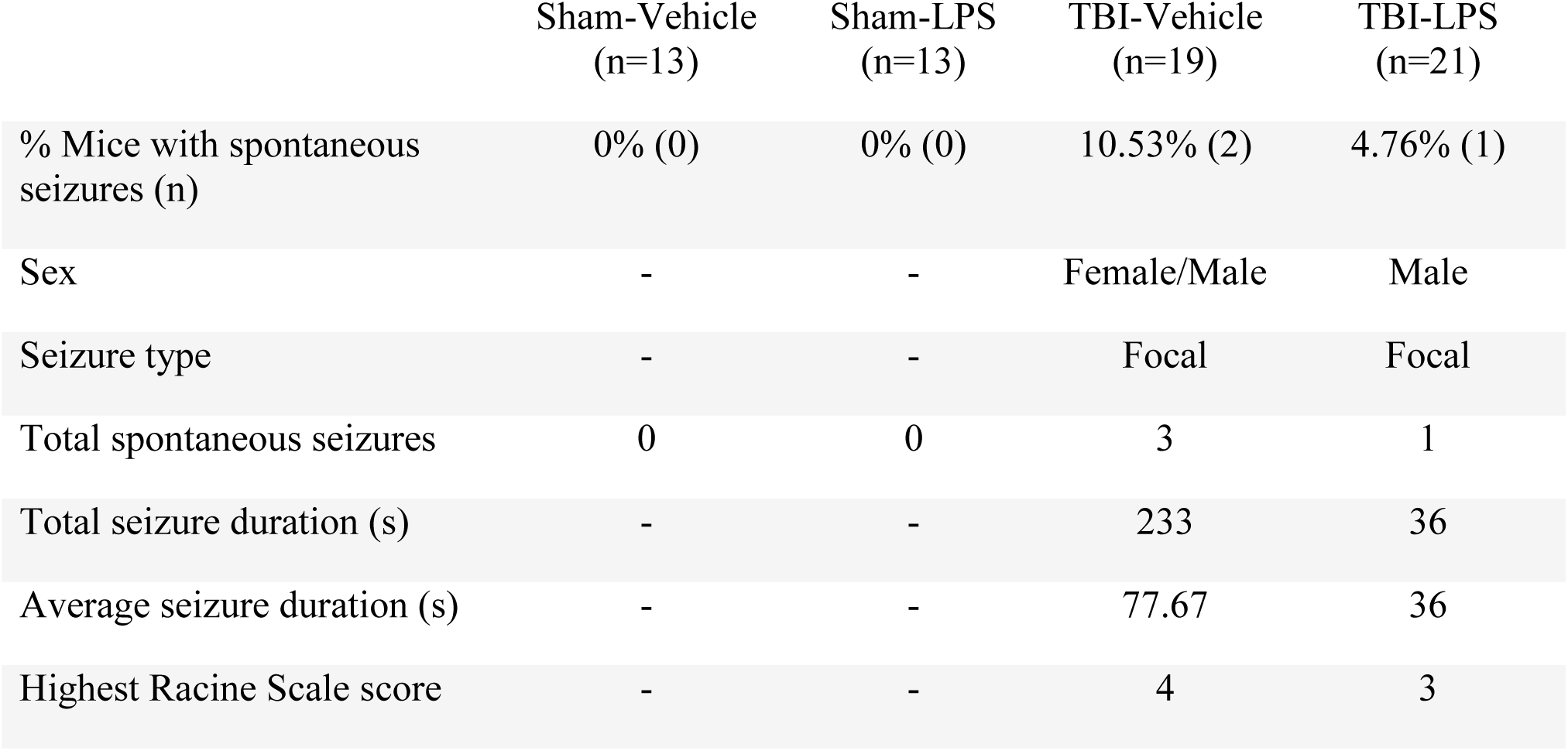
Spontaneous seizures detected by video-EEG monitoring chronically after TBI/LPS.

As a complementary or surrogate indicator of post-traumatic epilepsy, evoked seizure responses to low-convulsive dose PTZ were evaluated at the experimental end point of 24 weeks post-TBI. In response to PTZ administration, a significantly higher proportion of TBI-Vehicle and TBI-LPS mice displayed a convulsive seizure (90-91%) compared to Sham-Vehicle (47%) or Sham-LPS mice (33%) (Figure 6c; Chi-squared analysis p<0.0001). Consistently, TBI groups developed a convulsive seizure within a shorter time window (reduced latency; Figure 6d) compared to sham-operated groups (2-way ANOVA F_1,69_=38.88, p<0.0001). However, the duration of PTZ-evoked seizures was not different between experimental groups (Figure 6e).

Finally, the extent of neuropathology was assessed post-mortem by quantifying the extent of cortical and hippocampal tissue loss in the ipsilateral hemisphere, from cresyl violet-stained sections (Figure 7a-c). TBI resulted in a significant reduction in volume of both the cortex (F_1,19_=181.4, p<0.0001) and hippocampus (F_1,19_=18.25, p=0.0004). No effects of LPS were noted.

**Figure 7:**
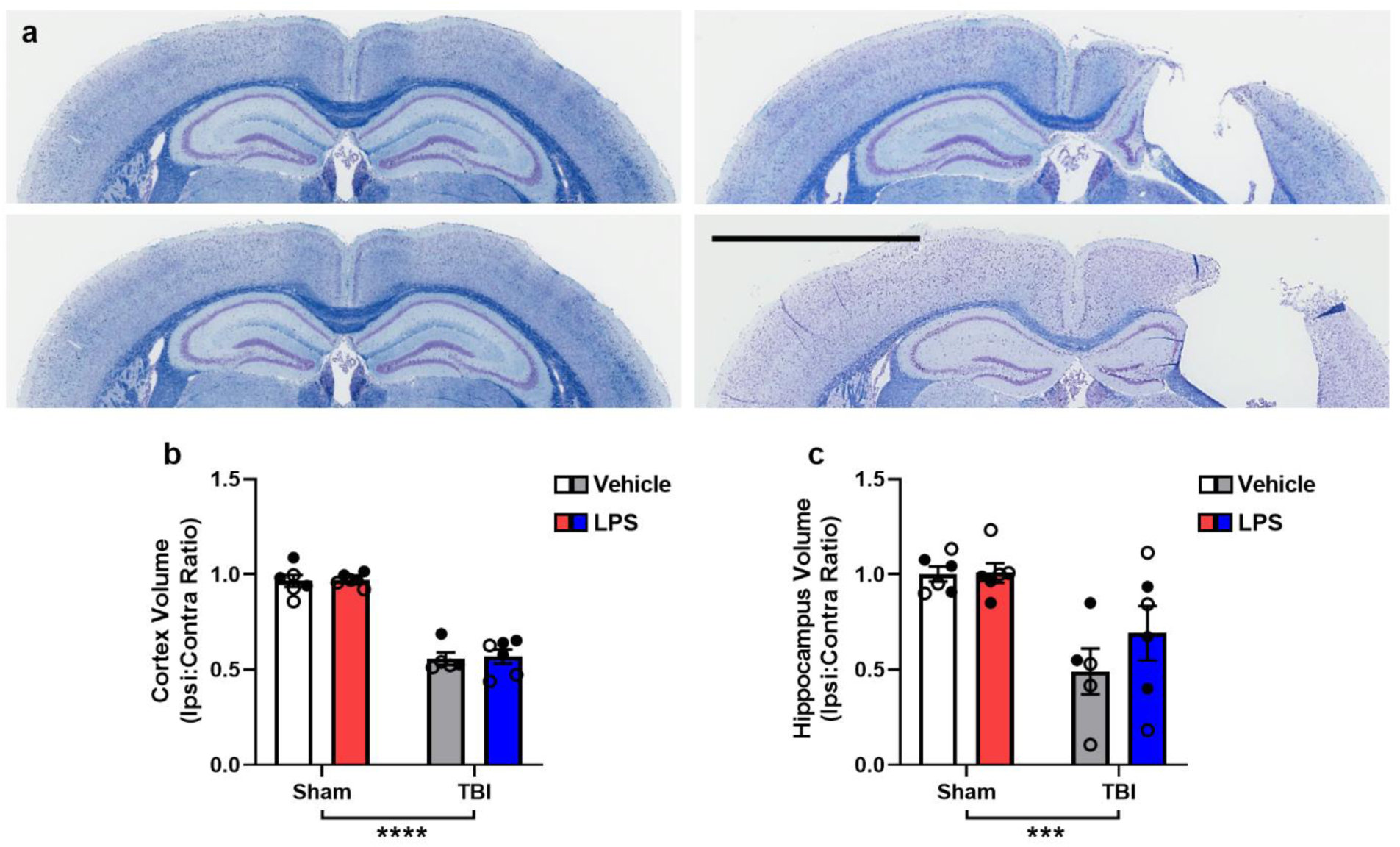
Chronic cortical and hippocampal atrophy in TBI mice was not affected by LPS. Representative figures of CV/LFB stained brain sections used for volumetric analysis from each group are depicted (a). Damage was measured as a ratio of ipsilateral to contralateral volume in the cortex and hippocampus. Two-way ANOVA observed an effect of TBI in both cortical (b) and hippocampal (c) ratio (****p < 0.0001; ***p = 0.0004), indicating atrophy in these regions chronically after TBI. Scale bar = 3 mm. n = 5-6/group.

## DISCUSSION

Patients acutely following a moderate-severe TBI are susceptible to nosocomial infections, presenting a significant challenge to an already-compromised immune system. However, the consequences and mechanisms by which this dual-insult worsens outcomes are poorly understood. In this study, we sought to determine the effects of LPS, as a systemic immune stimulus, on acute and chronic neurological and neurobehavioral outcomes following experimental moderate-severe TBI in young adult mice. As expected, LPS induced acute sickness behaviors including significant weight loss over the first two days post-administration, alongside transient hypoactivity and increased anxiety-like behavior.^1, 8^ Early systemic immune activation by LPS was confirmed by increased spleen weight and serum cytokines; yet TBI did not enhance these responses. In cortical and hippocampal brain tissue samples, gene expression analysis revealed a time course of inflammatory immune activation in the brains of injured or LPS-treated mice (e.g., IL-1β, IL-6, CCL2, TNFα), which was exacerbated in mice that received the combined insult of TBI and LPS. LPS-treated TBI mice also showed a reduction in fecal microbiome diversity at 24 h post-LPS, and changes in the relative abundance of key bacterial genera associated with sub-acute neurobehavioral and immune outcomes. Together, these acute post-injury findings indicate a complex interaction between the brain, immune system and gut microbiome. They also suggest that an exacerbated neuroinflammatory response and perturbation of the microbiome may be two of the mechanisms by which an additional immune challenge, such as a hospital-acquired infection, leads to health complications and poorer outcomes for patients.

Chronically, TBI induced long-term hyperactivity and cognitive deficits, brain atrophy, and increased seizure susceptibility by 6 months post-injury, consistent with previous studies using this experimental model.^8, 13, 22^ However, the additional insult of LPS administered early post-injury had no effect on these long-term outcomes. This suggests that the additional burden of LPS-induced inflammation resolved sub-acutely with no lasting effects on the evolution of the TBI neuropathology or neurological and neurobehavioral outcomes, including PTE.

Multiple lines of evidence point to inflammation as a key pathophysiological mechanism that underlies the development of post-traumatic epilepsy. For example, genetic variations in specific inflammatory genes (e.g., Il1β and its receptor IL-1R) have been associated with the risk post-traumatic epilepsy.^28^ Increased glial reactivity alongside the release of inflammatory mediators has been observed in the brain tissue of individuals with post-traumatic epilepsy, as well as in animal models of traumatic brain injury and epilepsy.^29-32^ Further studies have demonstrated effectiveness of anti-inflammatory pharmacological agents in reducing seizure incidence in animal models.^33, 34^ With this foundational understanding, we considered risk factors that an individual with TBI may face early after injury that could drive post-traumatic inflammation and epilepsy. Infections are an obvious candidate, being a known provocateur of a robust inflammatory response and a common comorbidity in patients with a TBI.^35-37^ In line with this hypothesis, we recently conducted a registry-based cohort study and reported an association between infections acquired during hospitalization after severe TBI and the risk of post-traumatic epilepsy at 2 years post-injury,^2^ suggesting that infections may promote epileptogenesis in the injured brain.

A frequent challenge when exploring this type of hypothesis in a preclinical setting is the low incidence of spontaneous seizures after experimental TBI. The incidence of post-traumatic epilepsy is variably reported from 4-50%, increasing with risk factors such as high injury severity, penetrating injury, and hemorrhage.^38-42^ In this CCI model of TBI, spontaneous seizures are most often reported in approximately 10% of experimental animals ^8, 22, 43, 44^ which is consistent with our current study findings. While some differences were seen in the duration, number and seizure focality, the low numbers prevented us from performing any statistical analysis and more robust conclusions about the effects of LPS on the development of PTE.. Similarly, while the PTZ challenge as a surrogate indicator of epileptogenesis indicated underlying hyperexcitability in the brains of TBI mice, consistent with previous studies,^8, 13, 22^ there was no additional impact of LPS on this response.

As a component of gram-negative bacterial cell walls, LPS is an endotoxin that is frequently used to mimic an infection-like host immune responses in *in vivo* model systems. We have previously used LPS derived from *Escherichia coli* as a tool to evaluate the effects of an infection-like immune challenge after experimental TBI in juvenile (post-natal day 21) mice.^8, 9^ In these past studies, we reported that the effects of TBI and LPS in juvenile mice were largely discrete and independent, both acutely and chronically, contrary to our expectations of interaction between these dual insults, but aligned with the findings observed herein. In the current study, we used young adult mice and LPS was derived from *Klebsiella pneumoniae*. *Klebsiella pneumoniae* is also a gram-negative bacterium and opportunistic pathogen that can cause serious infections in immunocompromised individuals, including pneumonia, urinary tract infections, bloodstream and wound infections.^45, 46^ Any differential findings between the current study and our previous findings may therefore be attributed to different ages of the model animal and/or different strains of LPS. While the timing of LPS administration relative to the injury has previously been shown to influence the subsequent immune response,^47^ in these studies LPS was administered at 4 days post-injury and intended to mimic an infection acquired within the first week or so post-injury, at a time when several aspects of the cellular inflammatory response are at their peak.^9, 48, 49^

The administration of isolated LPS via injection allows for tight control of the experimental paradigm. However, there are obviously several translational limitations of this model approach, including the administration route, use of a single bolus of endotoxin, and the fact that infectious agents are able to stimulate an immune response through multiple mechanisms beyond just LPS. We and others have called for a shift towards the use of live bacterial strains rather than isolated endotoxins or bacterial components, to better recapitulate the human scenario.^7^ Indeed, we have recently established a clinically-relevant model of experimental TBI combined with intratracheal administration of live *K. pneumoniae*, to replicate lung infection such as pneumonia in the first week after a brain injury.^10^ In this model, we observed acute sickness behaviors associated with a robust respiratory immune response following *K. pneumoniae*, and some interactions between *K. pneumoniae* and TBI in the brain in terms of inflammatory gene expression across an acute time course. However, surprisingly, no pronounced effects on the fecal microbiome were identified, perhaps due to the infection being largely localized to the lungs with more modest systemic effects compared to when LPS is administered i.p. Ongoing work in that model will evaluate the long-term consequences of this combined insult paradigm. Further, future studies should continue to optimize dual-insult models that incorporate both TBI and a secondary immune challenge, as this scenario is all-too-common for hospitalized patients and often complicates their recovery. Model systems can play an important role in dissecting the molecular mechanisms that drive poor outcomes, to reveal new therapeutic targets for pharmacological intervention.

Lastly, our findings provide novel insights into the complex interactions between the brain, immune system, and the gastrointestinal system. Increasing evidence demonstrates that TBI can impact the gut-brain axis and lead to imbalances in gut/fecal microbiota in terms of microbial composition and diversity.^25, 26, 50^ Such changes can have consequences for gut health and systemic inflammation, as well as exacerbate neurological symptoms and contribute to long-term recovery.^25, 26, 50^ Here, at the single time point of 24 h post-LPS/Vehicle (i.e., 5 days post-TBI/Sham), experimental TBI did not appear to alter the fecal microbiome. However, the combined insult of TBI+LPS resulted in a striking reduction in microbial diversity, including an increase in bacteria in the *Actinobacteria* and *Proteobacteria* phyla (e.g., *Proteus*) and decrease in bacterial genera such as *Prevotella*. Notably, levels of *Proteus* were negatively associated with several acute outcomes including levels of serum cytokines, inflammatory gene expression in the brain, and general activity of mice in an open field arena. These associations are intriguing, as *Proteus* bacteria are notoriously opportunistic and typically associated with gut dysbiosis and inflammation, with *P. mirabilis* found to be pathogenic in Crohn’s disease and ulcerative colitis.^51, 52^ Research by Choi and colleagues also suggests a link between *P. mirabilis* and neurological function – they found that *Proteus* isolated from a mouse model of Parkinson’s disease was able to induce motor deficits and reduced activity in the open field, alongside dopaminergic neuronal damage and neuroinflammation.^53^ Together, these findings warrant further exploration of the potential role of aberrant gut *Proteus* and other microbiome components in brain-gut-immune interactions in disease and injury contexts. Further, as our examination of the gut microbiome was limited to a single time point, future studies should consider the time course of such changes, as well as the effect of microbiome dysbiosis on gut structure and function.

## CONCLUSION

Clinical evidence indicates that infections acquired after TBI are associated with poorer outcomes, including an increased risk for post-traumatic epilepsy. Further work is required to optimize the experimental model paradigms to uncover the neurobiological mechanisms that underlie this relationship, and ensure the clinical relevance of preclinical study findings. The current study showed that an immune challenge early after experimental TBI, akin to a hospital-acquired infection, alters several aspects of the acute neuroinflammatory response to injury, although these are rapidly resolved. The chronic neurological and neurobiological consequences of TBI—including increased seizure susceptibility, memory deficits, and neuropathology—persisted for many months, and are not exacerbated by an LPS-mediated enhanced post-TBI immune response.

## Acknowledgements

The authors would like to thank the Monash Histology Platform, Monash Micro Imaging Platform, Monash Health Translation Precinct (MHTP) Medical Genomics Facility, Monash Animal Research Platform, and Crux Biolabs for their assistance with the project.

## Data Availability

Research datasets used in this study are available from the corresponding author upon reasonable request.

## Author Contributions

Project conceptualization: BDS, TOB, JL. Investigation, Methodology, Formal Analysis: BDS, AS, SSR, LC, MM, EC, NG, PCE. Resources: BDS, TOB, JL. Visualization: BDS. Writing – original draft: BDS, AS, SSR, MM. Writing – review and editing: All authors. We confirm that this manuscript is consistent with the Journal’s and publisher’s position on issues involved in ethical publication.

## REFERENCES

1. Sharma, R., Shultz, S.R., Robinson, M.J., Belli, A., Hibbs, M.L., O’Brien, T.J. and Semple, B.D. (2019). Infections after a traumatic brain injury: the complex interplay between the immune and neurological systems. Brain, Behavior, and Immunity 79, 63–74.

2. Chen, Z., Laing, J., Li, J., O’Brien, T.J., Gabbe, B.J. and Semple, B.D. (2024). Hospital-acquired infections as a risk factor for post-traumatic epilepsy: A registry-based cohort study. Epilepsia open 9, 1333–1344.

3. Shuman, E.K. and Chenoweth, C.E. (2018). Urinary Catheter-Associated Infections. Infectious disease clinics of North America 32, 885–897.

4. Vincent, J.L. (2003). Nosocomial infections in adult intensive-care units. Lancet 361, 2068–2077.

5. Zhang, X., Zhou, H., Shen, H. and Wang, M. (2022). Pulmonary infection in traumatic brain injury patients undergoing tracheostomy: predicators and nursing care. BMC pulmonary medicine 22, 130.

6. Clark, A., Zelmanovich, R., Vo, Q., Martinez, M., Nwafor, D.C. and Lucke-Wold, B. (2022). Inflammation and the role of infection: Complications and treatment options following neurotrauma. Journal of clinical neuroscience: official journal of the Neurosurgical Society of Australasia 100, 23–32.

7. Gandasasmita, N., Li, J., Loane, D.J. and Semple, B.D. (2023). Experimental Models of Hospital-Acquired Infections After Traumatic Brain Injury: Challenges and Opportunities. J Neurotrauma.

8. Sharma, R., Casillas-Espinosa, P.M., Dill, L.K., Rewell, S.S.J., Hudson, M.R., O’Brien, T.J., Shultz, S.R. and Semple, B.D. (2022). Pediatric traumatic brain injury and a subsequent transient immune challenge independently influenced chronic outcomes in male mice. Brain Behav Immun 100, 29–47.

9. Sharma, R., Zamani, A., Dill, L.K., Sun, M., Chu, E., Robinson, M.J., O’Brien, T.J., Shultz, S.R. and Semple, B.D. (2021). A systemic immune challenge to model hospital-acquired infections independently regulates immune responses after pediatric traumatic brain injury. J Neuroinflammation 18, 72.

10. Shad, A., Rewell, S.S.J., Macowan, M., Gandasasmita, N., Wang, J., Chen, K., Marsland, B., O’Brien, T.J., Li, J. and Semple, B.D. (2024). Modelling lung infection with Klebsiella pneumoniae after murine traumatic brain injury. J Neuroinflammation 21, 122.

11. Casillas-Espinosa, P.M., Andrade, P., Santana-Gomez, C., Paananen, T., Smith, G., Ali, I., Ciszek, R., Ndode-Ekane, X.E., Brady, R.D., Tohka, J., Hudson, M.R., Perucca, P., Braine, E.L., Immonen, R., Puhakka, N., Shultz, S.R., Jones, N.C., Staba, R.J., Pitkänen, A. and O’Brien, T.J. (2019). Harmonization of the pipeline for seizure detection to phenotype post-traumatic epilepsy in a preclinical multicenter study on post-traumatic epileptogenesis. Epilepsy Res 156, 106131.

12. Casillas-Espinosa, P.M., Sargsyan, A., Melkonian, D. and O’Brien, T.J. (2019). A universal automated tool for reliable detection of seizures in rodent models of acquired and genetic epilepsy. Epilepsia 60, 783–791.

13. Semple, B.D., O’Brien, T.J., Gimlin, K., Wright, D.K., Kim, S.E., Casillas-Espinosa, P.M., Webster, K.M., Petrou, S. and Noble-Haeusslein, L.J. (2017). Interleukin-1 Receptor in Seizure Susceptibility after Traumatic Injury to the Pediatric Brain. J Neurosci 37, 7864–7877.

14. Racine, R.J. (1972). Modification of seizure activity by electrical stimulation. II. Motor seizure. Electroencephalography and clinical neurophysiology 32, 281–294.

15. Clarke, E.L., Taylor, L.J., Zhao, C., Connell, A., Lee, J.J., Fett, B., Bushman, F.D. and Bittinger, K. (2019). Sunbeam: an extensible pipeline for analyzing metagenomic sequencing experiments. Microbiome 7, 46.

16. Bolger, A.M., Lohse, M. and Usadel, B. (2014). Trimmomatic: a flexible trimmer for Illumina sequence data. Bioinformatics (Oxford, England) 30, 2114–2120.

17. Wood, D.E., Lu, J. and Langmead, B. (2019). Improved metagenomic analysis with Kraken 2. Genome biology 20, 257.

18. McMurdie, P.J. and Holmes, S. (2013). phyloseq: an R package for reproducible interactive analysis and graphics of microbiome census data. PLoS One 8, e61217.

19. Paulson, J.N., Stine, O.C., Bravo, H.C. and Pop, M. (2013). Differential abundance analysis for microbial marker-gene surveys. Nature methods 10, 1200–1202.

20. Zamani, A., Powell, K.L., May, A. and Semple, B.D. (2020). Validation of reference genes for gene expression analysis following experimental traumatic brain injury in a pediatric mouse model. Brain Res Bull 156, 43–49.

21. Semple, B.D., Trivedi, A., Gimlin, K. and Noble-Haeusslein, L.J. (2015). Neutrophil elastase mediates acute pathogenesis and is a determinant of long-term behavioral recovery after traumatic injury to the immature brain. Neurobiol Dis 74, 263–280.

22. Webster, K.M., Shultz, S.R., Ozturk, E., Dill, L.K., Sun, M., Casillas-Espinosa, P.M., Jones, N.C., Crack, P.J., O’Brien, T.J. and Semple, B.D. (2019). Targeting high-mobility group box protein 1 (HMGB1) in pediatric traumatic brain injury: Chronic neuroinflammatory, behavioral, and epileptogenic consequences. Exp Neurol 320, 112979.

23. Ritchie, M.E., Phipson, B., Wu, D., Hu, Y., Law, C.W., Shi, W. and Smyth, G.K. (2015). limma powers differential expression analyses for RNA-sequencing and microarray studies. Nucleic acids research 43, e47.

24. Griffin, G.D. (2011). The injured brain: TBI, mTBI, the immune system, and infection: connecting the dots. Military medicine 176, 364–368.

25. Hanscom, M., Loane, D.J. and Shea-Donohue, T. (2021). Brain-gut axis dysfunction in the pathogenesis of traumatic brain injury. The Journal of clinical investigation 131.

26. Ma, E.L., Smith, A.D., Desai, N., Cheung, L., Hanscom, M., Stoica, B.A., Loane, D.J., Shea-Donohue, T. and Faden, A.I. (2017). Bidirectional brain-gut interactions and chronic pathological changes after traumatic brain injury in mice. Brain Behav Immun 66, 56–69.

27. Medel-Matus, J.S., Lagishetty, V., Santana-Gomez, C., Shin, D., Mowrey, W., Staba, R.J., Galanopoulou, A.S., Sankar, R., Jacobs, J.P. and Mazarati, A.M. (2022). Susceptibility to epilepsy after traumatic brain injury is associated with preexistent gut microbiome profile. Epilepsia 63, 1835–1848.

28. Diamond, M.L., Ritter, A.C., Failla, M.D., Boles, J.A., Conley, Y.P., Kochanek, P.M. and Wagner, A.K. (2015). IL-1beta associations with posttraumatic epilepsy development: A genetics and biomarker cohort study. Epilepsia 56, 991–1001.

29. Sharma, R., Leung, W.L., Zamani, A., O’Brien, T.J., Casillas Espinosa, P.M. and Semple, B.D. (2019). Neuroinflammation in Post-Traumatic Epilepsy: Pathophysiology and Tractable Therapeutic Targets. Brain sciences 9.

30. Webster, K.M., Sun, M., Crack, P., O’Brien, T.J., Shultz, S.R. and Semple, B.D. (2017). Inflammation in epileptogenesis after traumatic brain injury. J Neuroinflammation 14, 10.

31. Lucke-Wold, B.P., Nguyen, L., Turner, R.C., Logsdon, A.F., Chen, Y.W., Smith, K.E., Huber, J.D., Matsumoto, R., Rosen, C.L., Tucker, E.S. and Richter, E. (2015). Traumatic brain injury and epilepsy: Underlying mechanisms leading to seizure. Seizure 33, 13–23.

32. Salazar, A.M. and Grafman, J. (2015). Post-traumatic epilepsy: clinical clues to pathogenesis and paths to prevention. Handbook of clinical neurology 128, 525–538.

33. Saletti, P.G., Ali, I., Casillas-Espinosa, P.M., Semple, B.D., Lisgaras, C.P., Moshé, S.L. and Galanopoulou, A.S. (2019). In search of antiepileptogenic treatments for post-traumatic epilepsy. Neurobiol Dis 123, 86–99.

34. Dey, A., Kang, X., Qiu, J., Du, Y. and Jiang, J. (2016). Anti-Inflammatory Small Molecules To Treat Seizures and Epilepsy: From Bench to Bedside. Trends Pharmacol Sci 37, 463–484.

35. Kesinger, M.R., Kumar, R.G., Wagner, A.K., Puyana, J.C., Peitzman, A.P., Billiar, T.R. and Sperry, J.L. (2015). Hospital-acquired pneumonia is an independent predictor of poor global outcome in severe traumatic brain injury up to 5 years after discharge. J Trauma Acute Care Surg 78, 396–402.

36. Kumar, R.G., Kesinger, M.R., Juengst, S.B., Brooks, M.M., Fabio, A., Dams-O’Connor, K., Pugh, M.J., Sperry, J.L. and Wagner, A.K. (2020). Effects of hospital-acquired pneumonia on long-term recovery and hospital resource utilization following moderate to severe traumatic brain injury. J Trauma Acute Care Surg 88, 491–500.

37. Esnault, P., Nguyen, C., Bordes, J., D’Aranda, E., Montcriol, A., Contargyris, C., Cotte, J., Goutorbe, P., Joubert, C., Dagain, A., Boret, H. and Meaudre, E. (2017). Early-Onset Ventilator-Associated Pneumonia in Patients with Severe Traumatic Brain Injury: Incidence, Risk Factors, and Consequences in Cerebral Oxygenation and Outcome. Neurocritical care 27, 187–198.

38. Ferguson, P.L., Smith, G.M., Wannamaker, B.B., Thurman, D.J., Pickelsimer, E.E. and Selassie, A.W. (2010). A population-based study of risk of epilepsy after hospitalization for traumatic brain injury. Epilepsia 51, 891–898.

39. Frey, L.C. (2003). Epidemiology of posttraumatic epilepsy: a critical review. Epilepsia 44 Suppl 10, 11–17.

40. Pease, M., Gonzalez-Martinez, J., Puccio, A., Nwachuku, E., Castellano, J.F., Okonkwo, D.O. and Elmer, J. (2022). Risk Factors and Incidence of Epilepsy after Severe Traumatic Brain Injury. Ann Neurol 92, 663–669.

41. Ritter, A.C., Wagner, A.K., Fabio, A., Pugh, M.J., Walker, W.C., Szaflarski, J.P., Zafonte, R.D., Brown, A.W., Hammond, F.M., Bushnik, T., Johnson-Greene, D., Shea, T., Krellman, J.W., Rosenthal, J.A. and Dreer, L.E. (2016). Incidence and risk factors of posttraumatic seizures following traumatic brain injury: A Traumatic Brain Injury Model Systems Study. Epilepsia 57, 1968–1977.

42. Wang, X.P., Zhong, J., Lei, T., Wang, H.J., Zhu, L.N., Chu, S. and Liu, L. (2020). Epidemiology of traumatic brain injury-associated epilepsy in western China: An analysis of multicenter data. Epilepsy Res 164, 106354.

43. Bolkvadze, T. and Pitkanen, A. (2012). Development of post-traumatic epilepsy after controlled cortical impact and lateral fluid-percussion-induced brain injury in the mouse. J Neurotrauma 29, 789–812.

44. Statler, K.D., Scheerlinck, P., Pouliot, W., Hamilton, M., White, H.S. and Dudek, F.E. (2009). A potential model of pediatric posttraumatic epilepsy. Epilepsy Research 86, 221–223.

45. Keynan, Y. and Rubinstein, E. (2007). The changing face of Klebsiella pneumoniae infections in the community. International Journal of Antimicrobial Agents 30, 385–389.

46. Zembower, N.R., Zhu, A., Malczynski, M. and Qi, C. (2017). Klebsiella pneumoniae carbapenemase-producing K. pneumoniae (KPC-KP) in brain and spinal cord injury patients: potential for prolonged colonization. Spinal cord 55, 390–395.

47. Corrigan, F., Arulsamy, A., Collins-Praino, L.E., Holmes, J.L. and Vink, R. (2017). Toll like receptor 4 activation can be either detrimental or beneficial following mild repetitive traumatic brain injury depending on timing of activation. Brain Behav Immun 64, 124–139.

48. Hausmann, R., Kaiser, A., Lang, C., Bohnert, M. and Betz, P. (1999). A quantitative immunohistochemical study on the time-dependent course of acute inflammatory cellular response to human brain injury. Int J Legal Med 112, 227–232.

49. Jassam, Y.N., Izzy, S., Whalen, M., McGavern, D.B. and El Khoury, J. (2017). Neuroimmunology of Traumatic Brain Injury: Time for a Paradigm Shift. Neuron 95, 1246–1265.

50. He, Y., Wen, Q., Yao, F., Xu, D., Huang, Y. and Wang, J. (2017). Gut–lung axis: The microbial contributions and clinical implications. Critical reviews in microbiology 43, 81–95.

51. Zhang, J., Hoedt, E.C., Liu, Q., Berendsen, E., Teh, J.J., Hamilton, A., AW, O.B., Ching, J.Y.L., Wei, H., Yang, K., Xu, Z., Wong, S.H., Mak, J.W.Y., Sung, J.J.Y., Morrison, M., Yu, J., Kamm, M.A. and Ng, S.C. (2021). Elucidation of Proteus mirabilis as a Key Bacterium in Crohn’s Disease Inflammation. Gastroenterology 160, 317–330.e311.

52. Khorsand, B., Asadzadeh Aghdaei, H., Nazemalhosseini-Mojarad, E., Nadalian, B., Nadalian, B. and Houri, H. (2022). Overrepresentation of Enterobacteriaceae and Escherichia coli is the major gut microbiome signature in Crohn’s disease and ulcerative colitis; a comprehensive metagenomic analysis of IBDMDB datasets. Frontiers in cellular and infection microbiology 12, 1015890.

53. Choi, J.G., Kim, N., Ju, I.G., Eo, H., Lim, S.M., Jang, S.E., Kim, D.H. and Oh, M.S. (2018). Oral administration of Proteus mirabilis damages dopaminergic neurons and motor functions in mice. Scientific reports 8, 1275.

